# Structure and function of a 9.6 megadalton bacterial iron storage compartment

**DOI:** 10.1101/511345

**Authors:** T. W. Giessen, B. J. Orlando, A. A. Verdegaal, M. G. Chambers, J. Gardener, D. C. Bell, G. Birrane, M. Liao, P. A. Silver

**Affiliations:** Department of Systems Biology, Harvard Medical School, Boston, MA 02115, USA.; Wyss Institute for Biologically Inspired Engineering, Harvard University, Boston, MA 02115, USA.; Department of Biomedical Engineering, University of Michigan, Ann Arbor, MI 48109, USA.; Department of Cell Biology, Harvard Medical School, Boston, MA 02115, USA.; Center for Nanoscale Systems, Harvard University, Cambridge, MA 02138, USA.; School of Engineering and Applied Sciences, Harvard University, Cambridge, MA 02115,USA.; Department of Medicine, Beth Israel Deaconess Medical Center, Harvard Medical School, Boston, MA, 02215, USA.

## Abstract

Iron storage proteins are essential for maintaining intracellular iron homeostasis and redox balance. Iron is generally stored in a soluble and bioavailable form inside ferritin protein compartments. However, some organisms do not encode ferritins and thus rely on alternative storage strategies. Encapsulins, a class of protein-based organelles, have recently been implicated in microbial iron and redox metabolism. Here, we report the structural and mechanistic characterization of a 42 nm two-component encapsulin-based iron storage compartment from *Quasibacillus thermotolerans*. Using cryo-electron microscopy and x-ray crystallography, we reveal the assembly principles of a thermostable T = 4 shell topology and its catalytic ferroxidase cargo. We show that the cargo-loaded compartment has an exceptionally large iron storage capacity storing over 23,000 iron atoms. These results form the basis for understanding alternate microbial strategies for dealing with the essential element iron.

---

Iron is essential to all organisms on earth. However, the same properties that make iron useful for cellular metabolism can result in toxicity under aerobic conditions (1). Ferrous iron (Fe^2+^) is easily oxidized to insoluble ferric iron (Fe^3+^) resulting in the formation of harmful precipitates and reactive oxygen species (ROS)(2). Cells have evolved to cope with these problems by strictly controlling the intracellular concentration and reactivity of free iron (3). Ferritin proteins are used as the main iron storage system by animals, plants and most microbes (4). The main ferritin-like proteins involved in iron storage are ferritin (Ftn), bacterioferritin (Bfr) and DNA-binding proteins from starved cells (Dps) all able to oxidize Fe^2+^ to Fe^3+^ via a ferroxidase activity (5). While Ftn and Bfr are primarily used as a dynamic iron storage (6), the main function of Dps proteins is to counteract oxidative stress (7). Ferritins (Ftn and Bfr) assemble into 24 subunit protein compartments up to 12 nm in diameter able to store 2,000 to 4,000 Fe atoms in their interior (8, 9). However, some organisms do not encode ferritin genes in their genomes and their iron storage systems have remained elusive.

A newly discovered class of protein organelles called encapsulin nanocompartments are implicated in microbial iron and redox metabolism and have so far only been shown to be involved in oxidative stress response (10–13). Previously reported encapsulins share an HK97 phage-like fold and self-assemble from a single capsid protein into icosahedral compartments between 24 and 32 nm in diameter with triangulation numbers of T = 1 (60 subunits) and T = 3 (180 subunits), respectively (11, 12, 14). Their key feature is the ability to specifically encapsulate cargo proteins (Fig. 1A). Encapsulation is mediated by short C-terminal sequences referred to as targeting peptides (TPs) (12, 15). Genes encoding encapsulin shell proteins and dedicated cargo proteins are organized in co-regulated operons (10, 12). We have identified a novel type of encapsulin operon involved in iron metabolism in a range of Firmicutes we term the Iron-Mineralizing Encapsulin-Associated Firmicute (IMEF) system (10).

**Fig. 1.**
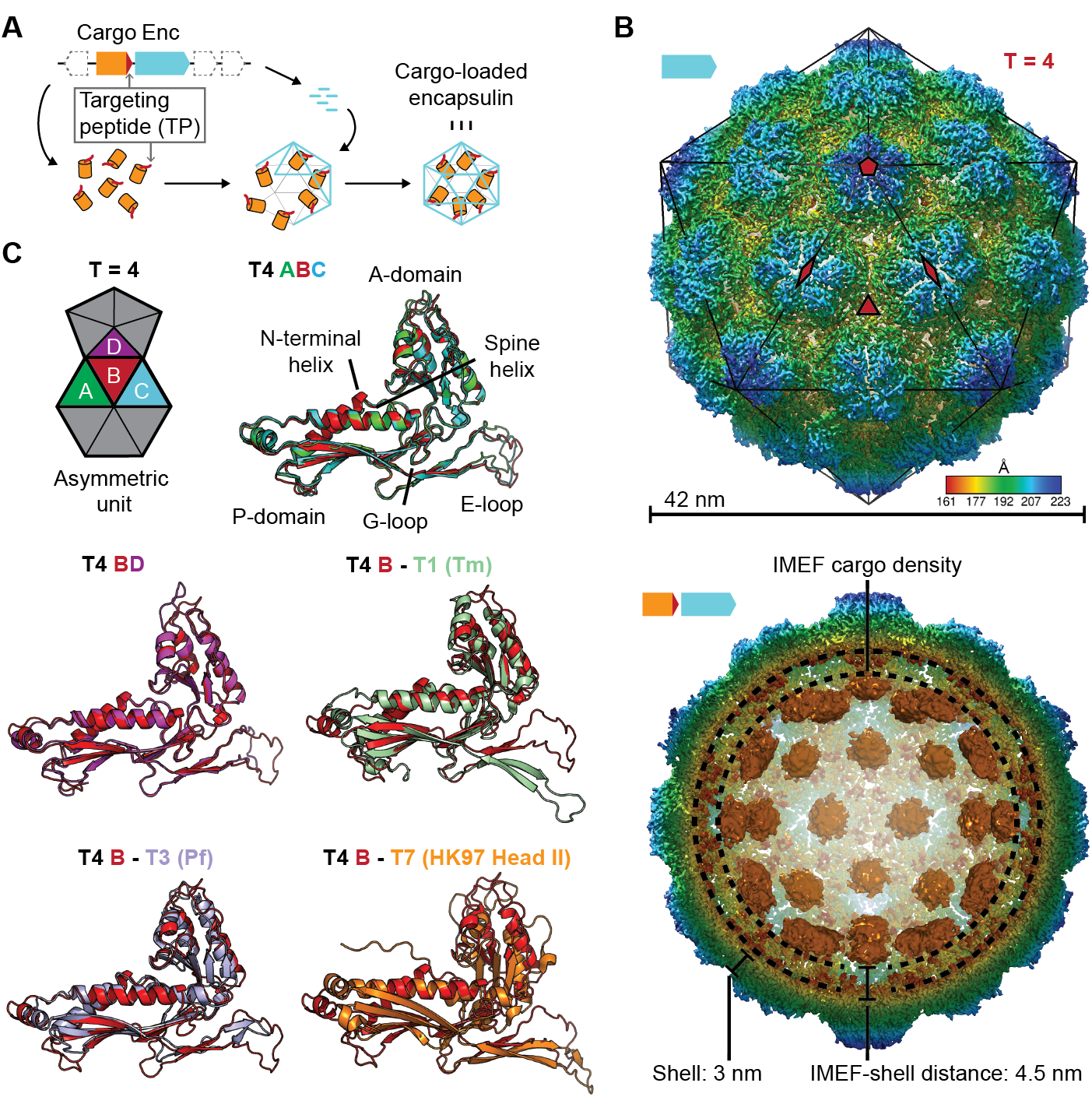
Overall architecture of the cargo-loaded T = 4 encapsulin. **(A)** Schematic diagram of a core encapsulin operon and targeting peptide (TP)-dependent cargo encapsulation. **(B)** Surface view of the cryo-EM map of the *Qs* T = 4 encapsulin shell (top) and inside view of cargo-loaded encapsulin (bottom). 5-, 3-and 2-fold symmetry axes are indicated by red symbols. The overall icosahedral symmetry is highlighted by black lines representing icosahedral facets. Cargo-densities are shown in orange while the shell is radially colored. To depict the complete cargo-loaded compartment, a 10 Å filtered map highlighting the cargo was combined with the 3.85 Å map of the shell. **(C)** Asymmetric unit of the T4 encapsulin shell and structural alignment of the four unique T4 shell monomers with one another and with the *T. maritima* (Tm) T = 1 monomer (3DKT), the *P. furiosus* (Pf) T = 3 monomer (2E0Z) and the HK97 bacteriophage Head II T = 7 monomer (2FT1).

Here, we report the structural and mechanistic characterization of the IMEF system found in *Quasibacillus thermotolerans (Qs)*, an organism that does not encode any ferritins in its genome. We show that this encapsulin-based system self-assembles into a thermostable 42 nm 9.6 MDa protein compartment with a novel T = 4 topology able to mineralize and store an exceptionally large quantity of iron.

IMEF systems are found in Firmicute genomes and their operon organization indicates a function in dynamic iron storage. We carried out BLASTp searches using IMEF cargo proteins as queries and identified 71 operons in a range of Firmicutes including *Qs* (Fig. S1). The core operon consists of the encapsulin capsid protein and the IMEF cargo protein with 70% of operons also encoding a 2Fe-2S ferredoxin homologous to bacterioferritin-associated ferredoxins (Bfd). Bfd proteins are involved in the mobilization of iron under iron-limited conditions (16). In addition, 31% of operons are associated with proteins similar to ferrochelatases involved in catalyzing the insertion of ferrous iron into protoporphyrins (17). The majority of IMEF-encoding genomes do not contain any ferritin or bacterioferritin genes (Table S1). Most IMEF genomes do however contain Dps-encoding genes. Overall, the operon organization of IMEF systems and the lack of other known primary iron storage proteins indicate a function for IMEF systems in dynamic iron storage similar to that of Ftn and Bfr.

Using a recombinant system, we produced homogeneous IMEF cargo-loaded encapsulins (Fig. S2). Through single-particle cryo-EM analysis, we determined the structure of the *Qs* IMEF encapsulin shell at an overall resolution of 3.85 Å (Fig. S3A). The IMEF encapsulin self-assembles into a 240 subunit icosahedral compartment with a diameter of 42 nm (Fig. 1B and Fig. S3A,B). The IMEF compartment is substantially larger than previously reported encapsulins and possesses a triangulation number of T = 4 instead of T = 1 or T = 3 (Fig. S4). The shell is composed of 12 pentameric and 30 hexameric capsomers occupying icosahedral vertices and faces, respectively. The 5-fold symmetry axes are located at the pentameric vertices while 3-fold symmetry axes are present at all interfaces where 3 hexameric capsomers meet. The center of each hexameric capsomer corresponds to an icosahedral edge possessing 2-fold symmetry. The icosahedral asymmetric unit consists of one pentameric and 3 hexameric monomers (Fig. 1B and Fig. S3C). Symmetrically arranged lower resolution density (ca. 10 Å) representing the IMEF cargo is visible in the compartment interior (Fig. 1B and Fig. S3D). 42 distinct densities, one for each capsomer of the T = 4 structure, can be observed. No connection of cargo and shell density is visible, likely due to the flexibility of the 37 amino acid linker preceding the IMEF targeting peptide that directs and anchors the IMEF cargo to the shell interior. The distance between the shell and cargo densities is 4.5 nm which can be bridged by the 37 amino acid linker.

The 4 capsid proteins of the asymmetric unit adopt different conformations with significant differences found in the E-loop and A-domain (Fig. 1C). E-loops are located at capsomer interfaces and their relative orientation plays a key role in determining the overall topology and triangulation number of encapsulin compartments as evidenced by comparison of the IMEF T = 4 monomer with T = 1, T = 3 and T = 7 capsid proteins. A-domain loops form compartment pores and are likely adapted to the particular function of a given encapsulin explaining the observed conformational diversity. In addition, local resolution maps indicate that E-loops and A-domain loops represent the most flexible parts of the shell (Fig. S5).

The IMEF compartment possesses a non-covalent chainmail topology and is highly thermostable. E-loops and P-domains of neighboring capsid monomers arrange head to tail to form interlocking concatenated rings resulting in a non-covalent chainmail topology (Fig. 2A) (18). In contrast to HK97 where an isopeptide bond covalently links E-loops and P-domains (19), the IMEF encapsulin uses non-covalent interactions. At each 3-fold pore, E-loops connect with two neighboring P-domains including the G-loop conserved in T = 4 encapsulins and their interfaces contain complementary electrostatic as well as aromatic and potential anion-π interactions (Fig. 2B and Fig. S6) (20). The IMEF cargo and shell protein are both highly thermostable with melting temperatures of 80.6 and 86.6°C, respectively (Fig. 2C). A stabilizing effect is observed for the cargo-loaded compartment (88.9°C). Compartments isolated from high iron conditions show even greater thermal stability (91.8°C) likely due to the internal cavity being stabilized by mineralized material.

**Fig. 2.**
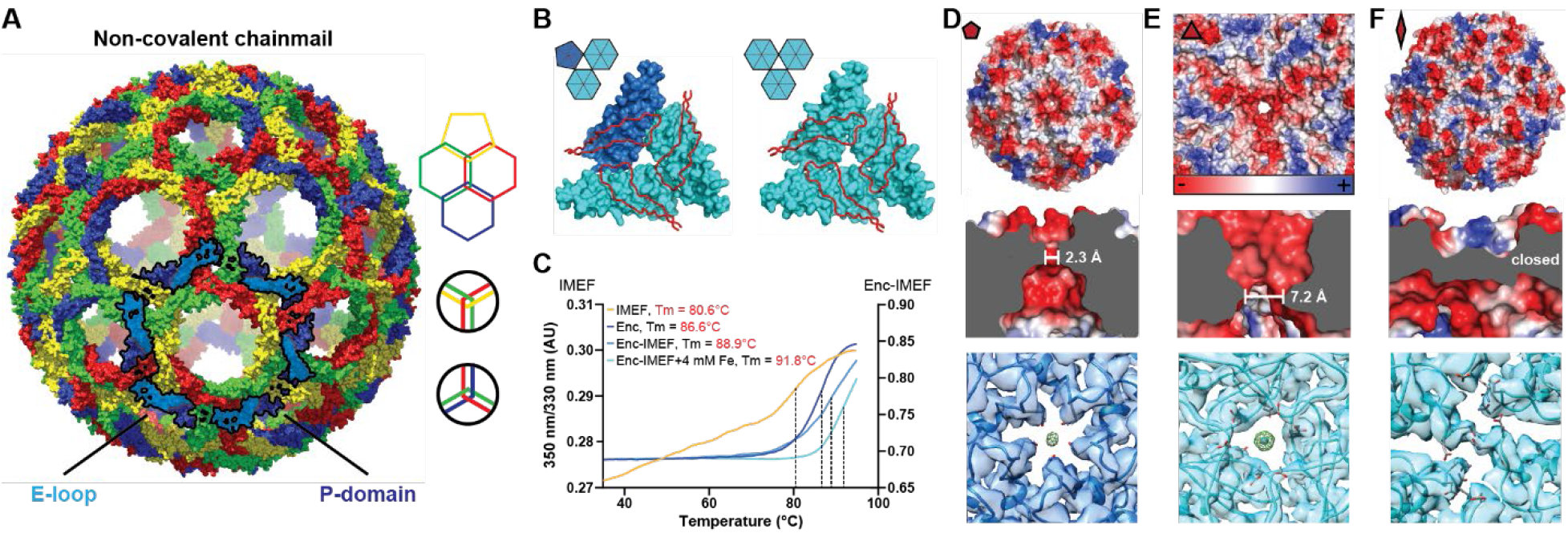
Non-covalent chainmail topology, thermal stability and pores of the T = 4 encapsulin shell. **(A)** Chainmail network mediated by E-loop and P-domain interactions. Only E-loops and P-domains are shown. E-loops and P-domains of the outlined ring belonging to the same monomer are located next to one another and are shown in light and dark blue, respectively. **(B)** Extended E-loop interactions interlock neighboring capsid monomers at the two unique threefold interfaces. Each E-loop interacts with two P-domains. **(C)** Representative thermal unfolding curves for *Qs* T = 4 encapsulin components determined via differential scanning fluorimetry. Tm: midpoint of the thermal unfolding curve. (**D, E** and **F**) Electrostatic surface representation of the 5-fold (D) and 3-fold (E) T = 4 shell pores and the 2-fold symmetry axis (F). Outside views showing negatively charged pores (top) with no pore opening observed at the two-fold symmetry axis, cutaway side view highlighting the narrowest point of the pores (middle) and cryo-EM maps with fitted monomer models in ribbon representation (bottom). Additional cryo-EM density is observed at the center of both pores in interaction distance with the side chains of pore residues (5-fold: Asn200, 3-fold: Asp9, Asp71, Glu251 and Glu252, shown in stick representation).

The IMEF encapsulin shell contains negatively charged pores at the 3- and 5-fold symmetry axes. The surface view of the intact shell (Fig. S7) shows a tight packing with pores at the 3- and 5-fold symmetry axes and at the interface between two hexameric and one pentameric capsomer (pseudo 3-fold) representing the only conduits to the interior. All pores are negatively charged on both the exterior and interior surface due to the presence of conserved aspartate, glutamate and asparagine residues (Fig. 2D, 2E, Fig. S8 and Fig. S9). The 3-fold pore forms the largest channel to the IMEF compartment interior and is 7.2 Å wide at its narrowest point, substantially larger than previously reported encapsulin pores (Fig. S10). Extra cryo-EM density is observed at the center of both the 3-fold and 5-fold pores potentially resulting from bound positively charged ions. The 2-fold symmetry axes at the center of hexameric capsomers also represent potential channels, as observed in T = 3 systems (21), but the conformation of two asparagine side chains prevents the formation of a 2-fold opening in the T = 4 shell leading to a closed pore (Fig. 2F).

This observation combined with the flexibility observed for loops around the 2- and 5-fold symmetry axes in local resolution maps (Fig. S5) indicates gated pores in encapsulins that may regulate ion flux to the compartment interior, similar to some ferritins (22).

Sequence and x-ray structure analysis show that the IMEF cargo represents a distinct class of ferritin-like protein (Flp) with an unusual ferroxidase center. Phylogenetic analysis revealed that the IMEF cargo protein is a member of the Flp superfamily and is most closely related to Dps proteins (Fig. 3A and Table S3) but no known ferroxidase motifs could be detected at the sequence level (5). All IMEF proteins share a conserved C-terminal TP (Fig. 3B). We determined the x-ray crystal structure of the IMEF cargo to a final resolution of 1.72 Å (Fig. 3C and Table S4). The cargo adopts a four-helix bundle fold characteristic of other members of the Flp superfamily and forms a dimer with two Fe atoms bound at the subunit interface creating a ferroxidase site based on an alternative ferroxidase sequence motif (Fig. 3D, Fig. S11 and Fig. S12). This leads to a combined molecular weight of the fully cargo-loaded IMEF compartment of 9.6 MDa (42 × cargo dimer [22.6 kDa] + 240 × capsid protein, [32.2 kDa]). Removal of the 13 C-terminal residues results in empty encapsulin shells confirming that the IMEF TP is necessary for cargo encapsulation (Fig. 3E).

**Fig. 3.**
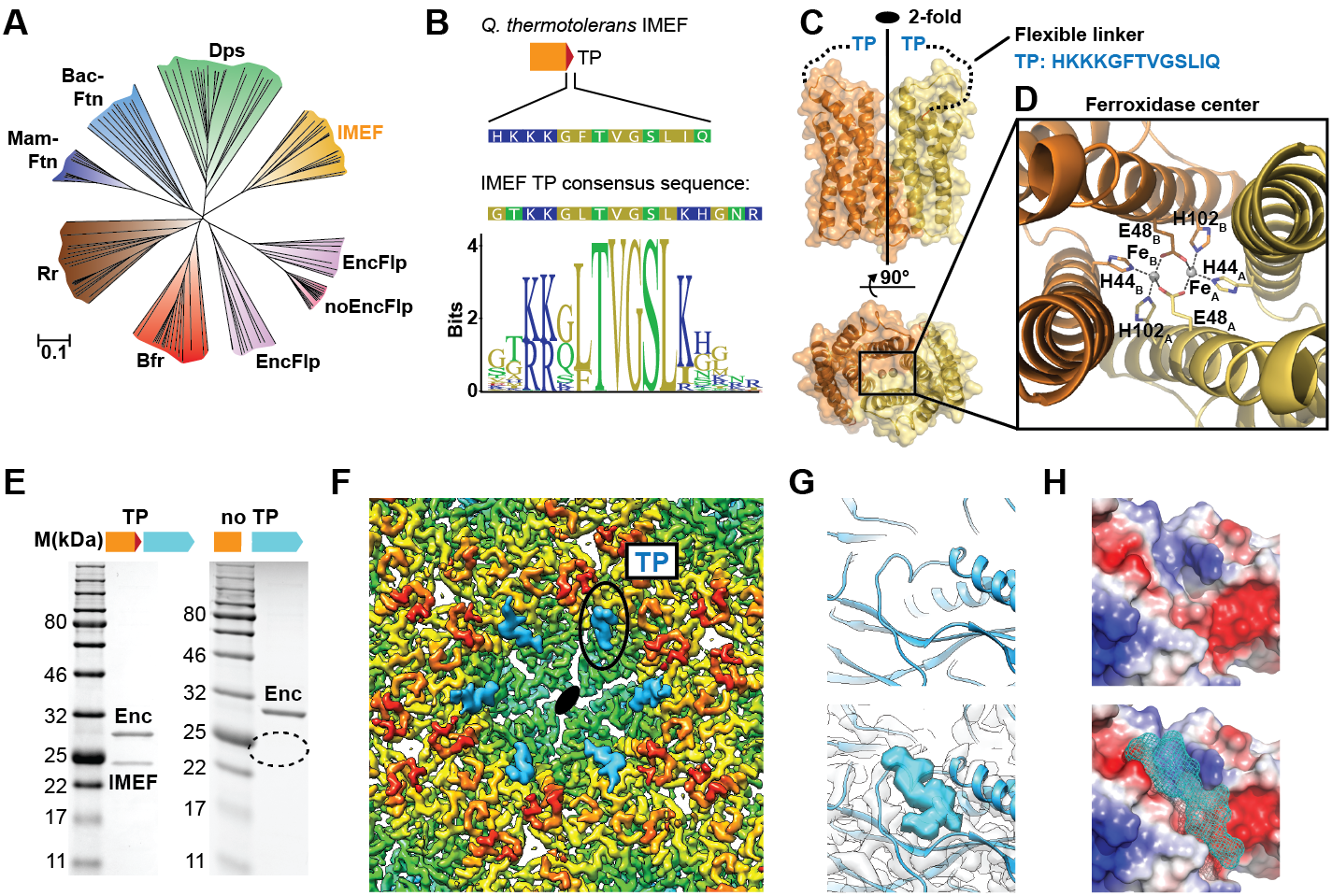
Structure and analysis of the IMEF cargo protein and TP-mediated cargo-shell coassembly. **(A)** Neighbor-joining phylogeny (cladogram) of protein classes involved in iron metabolism that are part of the Flp superfamily. Scale bar: amino acid substitutions per site. EncFlp: Flps found within encapsulin operons containing TPs, noEncFlp: Flps found outside encapsulin operons not containing TPs, Rr: rubrerythrins, Mam-Ftn: mammalian ferritins, Bac-Ftn: bacterial ferritins. **(B)** TP sequence of the *Qs* IMEF cargo protein and TP sequence logo highlighting strong sequence conservation. **(C)** X-ray crystal structure of the *Qs* IMEF cargo. **(D)** Di-iron ferroxidase active site of the IMEF cargo. The iron-coordinating residues are shown in stick representation. **(E)** SDS-PAGE gels of purified encapsulins showing that co-purification is dependent on the presence of the TP. **(F)** Cryo-EM map interior view of the 2-fold symmetry axis with TP density shown in cyan. **(G)** Close-up of additional cryo-EM density observed around the 2-fold symmetry axis. **(H)** Electrostatic surface representation of the TP binding site without (top) and with (bottom) TP. The 7 C-terminal IMEF residues are shown as a surface mesh.

Additional cryo-EM density around the 2- and 5-fold symmetry axes reveals TP-binding sites and illuminates cargo-shell co-assembly. Through analysis of the T = 4 cryo-EM map, additional densities were identified that could not be explained by the encapsulin capsid protein (Fig. 3F). These densities represent bound TPs anchoring IMEF cargo to the interior surface of the compartment. Even though only 42 cargo densities are observed, TP densities can be found at all 240 capsid monomers indicating averaging during cryo-EM reconstruction. Strong TP density is observed for all 180 monomers that are part of 2-fold symmetrical hexameric capsomers (Fig. 3F) while substantially weaker density is found for TPs bound to the 60 pentameric monomers (Fig. S13A) thus revealing higher occupancy and preferential TP binding around 2-fold symmetry axes which can be explained by different binding site conformations (Fig. S13) and higher local shell mobility (Fig. S5). The main TP binding sites surrounding the 2-fold symmetry axes are formed by conserved residues of the P-domain and N-terminal helix (Fig. S9) similar to the *T. maritima* T = 1 encapsulin system (12). Thus, the presence of the N-terminal helix and the resulting binding site generally underpin encapsulins’ ability to interact with TPs and encapsulate cargo proteins. The TP residues TVGSLIQ were tentatively built and refined into the additional density present at hexameric capsomers producing a model with good geometry (Fig. S13C). The TP binds to a surface groove based on shape complementarity and two key ionic interactions with highly conserved positively charged residues locking the TP in place (Fig. S13C).

Heterologous expression of the IMEF core operon in *E. coli* leads to *in vivo* formation of large Fe- and P-rich electron-dense particles. Thin section negative stain transmission electron microscopy (TEM) of *E. coli* cells grown in Fe-rich (4 mM) medium and expressing the *Qs* IMEF core operon results in the formation of clusters of large intracellular electron-dense particles (Fig. 4A and Fig. S14). Scanning TEM-energy-dispersive x-ray spectroscopy (EDS) revealed that these particles primarily contain uniformly distributed Fe, P and O with an estimated Fe:P ratio near 1 (Fig. 4B). Selected area electron diffraction (SAED) further indicates that this mineralized material is amorphous (Fig. S15), similar to bacterioferritin systems (23).

**Fig. 4.**
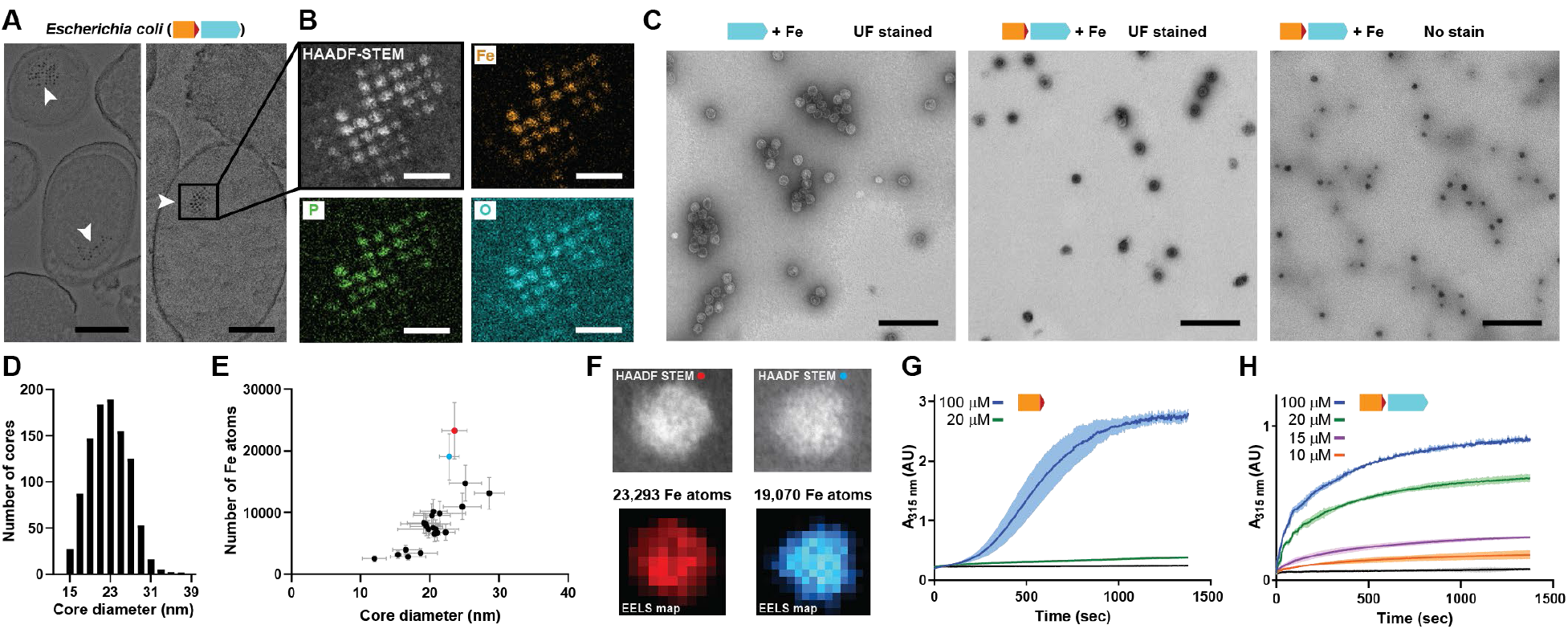
Mineralization of large iron-rich particles by the T = 4 encapsulin. **(A)** Thin section micrographs of *E. coli* heterologously expressing the *Qs* IMEF core operon. Electron-dense particles often cluster together in regular arrays. Scale bars: 500 nm (left), 400 nm (right). **(B)** Close-up high angle angular dark field (HAADF) scanning TEM and EDS maps of a cluster of particles showing Fe, P and O as the main particle constituents. Scale bars: 100 nm. **(C)** Micrographs of uranyl formate (UF)-stained encapsulins produced in and isolated from *E. coli*grown in high iron media expressing the capsid protein alone (left) or the core operon (middle and right). Without UF stain, electron-dense particles are clearly visible (right). Scale bars: 250 nm. **(D)** Size distribution of electron-dense particles in unstained micrographs. **(E)** Electron energy loss spectroscopy (EELS) of 22 select cores. **(F)** HAADF-STEM micrographs and EELS maps of the two highlighted cores from (E). **(G)** *In vitro* ferroxidase assay of purified IMEF cargo at different Fe^2+^ concentrations. Mean values resulting from technical triplicates and error bands using standard deviation are shown. **(H)** Ferroxidase assay of cargo-loaded T = 4 encapsulin at different Fe^2+^ concentrations.

The IMEF encapsulin mineralizes up to 30 nm Fe-rich cores in its interior with up to 23,000 Fe atoms stored per particle. IMEF encapsulins purified from *E. coli* grown under high Fe conditions contain electron dense cores visible in unstained samples with an average diameter of 23 nm (Fig. 4C,D and Fig. S16). The largest observed particles are up to 30 nm in diameter. The theoretical size limit imposed by the T = 4 encapsulin protein shell is 36 nm and particles close to this size are observed in thin-sections of *Geobacillus* natively encoding the IMEF system (Fig. S17). EDS analysis of particles isolated from *E. coli* and comparison with standards indicate a very similar elemental composition and elemental distribution as observed for thin section samples with a Fe:P ratio of 1:1.1 (Fig. S18). To determine the number of iron atoms stored per particle, we carried out electron energy loss spectroscopy (EELS) on purified Fe-loaded compartments (Fig. 4E,F and Fig. S19). The highest observed number of stored Fe per particle was 23,293 (23.6 nm) (Table S5). Extrapolating to the maximum theoretical particle diameter of 36 nm and the highest density observed (3.40 Fe atoms/nm^3^) leads to a maximum number of Fe atoms that can be stored by the IMEF system of around 83,000 (Table S5).

To learn more about the mechanism of iron mineralization, we assayed peroxidase and ferroxidase activity using H_2_O_2_ and O_2_ as the oxidants, respectively. No peroxidase activity was detected. For the IMEF cargo alone, a sigmoidal ferroxidase iron oxidation curve was observed indicative of autocatalytic Fe oxidation taking place at newly formed mineral surfaces (24, 25). However, assaying the cargo-loaded encapsulin results in a typical hyperbolic enzyme catalysis curve. These observations imply that the encapsulin shell controls the flux of iron to the interior of the compartment thus preventing uncontrolled autocatalytic mineralization (Fig. S20).

Our structural model and functional analysis of the IMEF encapsulin system reveal an alternative way to store large amounts of Fe independent of ferritins. The IMEF system can in principle store more than 20 times more Fe than Ftn or Bfr systems. In contrast to ferritin systems, IMEF encapsulins are two-component systems with the catalytic activity separated from the protein shell. The IMEF cargo protein is flexibly tethered and primarily localizes 4.5 nm away from the capsid interior. This suggests that once iron has been channeled to the encapsulin interior via pores, it diffuses to the ferroxidase active site of the IMEF cargo, making it necessary to strictly control interior iron concentration to prevent runaway mineralization. In sum, we have elucidated the structure and mechanism of the largest iron storage complex to date indicating that alternative systems exist across nature to address the critical problem of safe and dynamic iron storage.

## Acknowledgments

We thank M. Ericsson, P. Coughlin and L. Trakimas for technical support (HMS Electron Microscopy Facility).

## Authors contributions

T.W.G. and P.A.S. conceived and designed the study. T.W.G. carried out all cloning and performed biochemical and negative staining TEM experiments. B.J.O. and M. L. performed cryo-EM image processing, data analysis and model building. M.G.C. collected cryo-EM data sets. A.A.V. purified and crystalized the IMEF cargo protein and cultivated *Geobacillus*. J.G. and D.C.B. performed and analyzed EDS and EELS experiments. G.B. helped in crystallization experiments and solved the IMEF x-ray structure. T.W.G. and P.A.S. wrote the manuscript. B.J.O. and M.L. helped in editing the manuscript.

## Data and materials availability

A cryo-EM density map of the cargo-loaded IMEF encapsulin has been deposited in the Electron Microscopy Data Bank under the accession number 9383. The corresponding atomic coordinates for the atomic model have been deposited in the Protein Data Bank (accession number: 6NJ8). Atomic coordinates for the IMEF cargo protein have been deposited in the Protein Data Bank under accession number 6N63.

Correspondence and requests for materials should be addressed to the corresponding authors.

## Supplementary Materials

Materials and Methods

Fig. S1 – S20

Table S1 – S5

References (26 – 49)

