## Supplementary Material for "Structure and function of a 9.6 megadalton bacterial iron storage compartment"

**This PDF file includes:**

Materials and Methods
Figs. S1 to S20
Tables S1 to S5
References (26 – 49)

### Materials and Methods

Computational analysis of genomes, encapsulin gene clusters, sequences and protein structures

Initial identification of IMEF systems was achieved by utilizing the Enzyme Similarity Tool (ESI) in combination with the Genome Neighborhood Network Tool (GNT) of the Enzyme Function Initiative (EFI) (*26*). The previously identified IMEF cargo protein from *Q. thermotolerans* (WP_039238471) was used as a query to initiate an ESI Sequence BLAST search of the UniProt database. UniProt BLAST Query E-value was chosen to be 5. After the initial dataset was created, we used an alignment score (based on the alignment score vs percent identity plot) that would correspond to a percent identity of 20 for initial outputting and interpretation of protein sequences and sequence similarity networks (SSNs). The resulting xgmml network file was then submitted to GNT. The resulting Genome Neighborhood Diagrams of all identified IMEF operons where analyzed using the GNT diagram explorer and operon diagrams were downloaded as svg files.

Genomes of IMEF system-encoding organisms were searched for Ftn, Bfr and Dps proteins using NCBI's blastp suite. As queries, Firmicute homologs of ferritin, bacterioferritin and Dps were used (Ftn: OTY20392, Bfr: EEK74551, Dps: WP_039234032).

Phylogenetic analysis was based on Clustal Omega (ClustalO) alignments carried out using the default settings of the Multiple Sequence Alignment online tool of the European Molecular Biology Laboratory's European Bioinformatics Institute (EMBL-EBI). A nearest-neighbor phylogenetic trees based on the ClustalO alignment were generated using the Simple Phylogeny Tool at EMBL-EBI. Alignments and trees were then annotated and analyzed using Geneious 9.1.4.

Cryo-EM data and structural models were analyzed using UCSF Chimera 1.13.1rc and Open Source PyMOL 1.8.x. Structural alignments of capsid protein monomers were carried out in PyMOL using the align command. The IMEF model used for molecular replacement was generated using the I-TASSER webserver (*27, 28*).

Molecular biology and cloning

All constructs used in this study were ordered as gBlock Gene Fragments from Integrated DNA Technologies (IDT). Codon usage was optimized for *E. coli* expression using the IDT Codon Optimization Tool with the amino acid sequences of the respective proteins of interest as input. For the IMEF operon containing multiple genes, intergenic regions were not changed. The IMEF cargo protein construct was ordered with a C-terminal $His_6$ tag. For the operon construct containing the TP-less IMEF cargo, the 13 C-terminal residues (HKKKGFTVGSLIQ) were omitted from the IMEF cargo protein, thus removing the TP.

Gibson Assembly® Master Mix was obtained from New England BioLabs (NEB). DNA sequencing was carried out by GENEWIZ. MegaX DH10B T1R electrocompetent *E. coli* cells (ThermoFisher) were used for all cloning procedures while One Shot® BL21 Star (DE3) chemically competent *E. coli* cells (Invitrogen) were used for protein production and all other experiments. pETDuet1 was used as the expression vector for all constructs. For the construction of expression vectors, Gibson Assembly was employed. gBlock Gene Fragments containing 20 bp overlaps for direct assembly were combined with NdeI and PacI digested pETDuet1 resulting in assembled expression vectors (fragments were inserted in MCS2). Electrocompetent *E. coli* DH10B cells were transformed and the resulting plasmids confirmed via sequencing.

Expression and purification of proteins and protein compartments

All non-high iron expression experiments were carried out in lysogeny broth (LB) supplemented with ampicillin (100 μg/mL). Size exclusion chromatography/gel filtration for capsid purification was performed with an ÄKTA Explorer 10 (GE Healthcare Life Sciences) equipped with a HiPrep 16/60 Sephacryl S-500 HR column (GE Healthcare Life Sciences). For analytical size exclusion, a Superdex 200 10/300 GL column (GE Healthcare Life Sciences) was used. Protein samples were concentrated using Amicon Ultra Filters (Millipore). For SDS-PAGE analysis, 14% Novex Tris-Glycine Gels (ThermoFisher Scientific) were used. DNA concentrations were measured using a Nanodrop ND-1000 instrument (PEQLab).

Sequence-confirmed plasmids were used to transform *E. coli* BL21 (DE3) Star cells (0.5 ng total plasmid DNA). Resulting colonies were used to inoculate pre-expression cultures.

For large scale protein expressions, 500 mL of LB in 2 L baffled flasks were inoculated (1:50) using an over-night culture, grown at 37°C and 200 rpm to an $OD_{600}$ of 0.5. The temperature was then shifted to 30°C and the cultures induced with IPTG (final concentration: 0.05 mM). Cultures were grown at 30°C for 18 h, harvested through centrifugation (4000 rpm, 15 min, 4°C) and pellets either immediately used or frozen in liquid nitrogen and stored at -20°C for later use.

For encapsulin and His-tagged protein purifications, pellets were thawed, resuspended in 5 mL Tris buffer (50 mM Tris, 150 mM NaCl, pH 8), then lysozyme (1 mg/mL) and DNaseI (1 μg/mL) were added and the cells incubated on ice for 20 min. Cell suspensions were subjected to sonication using a 550 Sonic Dismembrator (FisherScientific). Power level 3.25 was used with a pulse time of 8 sec and an interval of 10 sec. Total pulse time was 4 min. Cell debris was subsequently removed through centrifugation (8000 rpm, 15 min, 4°C). The cleared supernatant was then used either for protein affinity or encapsulin compartment purification.

His-tagged IMEF cargo was purified using Ni-NTA agarose resin (Qiagen) via the batch Ni-NTA affinity procedure following the supplier's instructions. Buffer A (50 mM Tris, 150 mM NaCl, 20 mM imidazole, pH 8) was used to wash the resin after protein binding and buffer B (50 mM Tris, 150 mM NaCl, 250 mM imidazole, pH 8) was used to elute bound protein. Samples were concentrated and dialyzed using Amicon filters (10 kDa molecular weight cutoff) and Tris (pH 7.4) buffer and evaluated using SDS-PAGE. Further analyses were carried out directly or the next day with protein being stored on ice.

For encapsulin purification, 0.1 g NaCl and 0.5 g of PEG-8000 were added (10% w/v final concentration) to 5 mL cleared lysate, followed by incubation on ice for 20 min. The precipitate was collected through centrifugation (8000 rpm, 15 min, 4°C), suspended in 3 mL Tris (pH 8) buffer and filtered using a 0.2 μm syringe filter. The samples were then subjected to size exclusion chromatography using Tris (pH 8) buffer and a flow rate of 1 mL/min.

Fractions were evaluated using SDS-PAGE analysis and encapsulin-containing fractions were combined, concentrated and dialyzed using Amicon filters (100 kDa molecular weight cutoff) and Tris buffer without NaCl (20 mM Tris, pH 8).

The low salt sample was then loaded on a HiPrep DEAE FF 16/10 Ion Exchange column (GE Healthcare Life Sciences). The gradient used for ion-exchange chromatography was as follows: 100% A for 0-100 mL, 100% A to 50% A + 50% B for 100-200 mL, 100% B for 200-300 mL, 100% A for 300-400 mL (A: 20 mM Tris, pH 8, B: 20 mM Tris, 1 M NaCl, pH 8, flow rate: 3 mL/min). Again, SDS-PAGE was used to identify product fractions followed by Amicon filter concentration and buffer exchange to Tris buffer (50 mM Tris, 150 mM NaCl, pH 8).

Final samples were either directly subjected to additional experiments or stored on ice overnight.

<u>Negative stain transmission electron microscopy (TEM) of purified encapsulins</u>

200 Mesh Gold Grids (FCF-200-Au, EMS) were used for all negative stain TEM experiments. TEM experiments of negatively stained protein samples were carried out at the HMS Electron Microscopy Facility using a Tecnai G2 Spirit BioTWIN instrument.

For negative-staining TEM, encapsulin samples were diluted to 1-10 μM using Tris buffer (50 mM Tris, 150 mM NaCl, pH 8) and subsequently adsorbed onto formvar/carbon coated gold grids. Prior to applying 5 μL of diluted sample, grids were glow-discharged using a 100x glow discharge unit (EMS) to increase their hydrophilicity (10 sec, 25 mA). After 1 min adsorption time, excess liquid was blotted off using Whatman #1 filter paper, washed one time with distilled $H_2O$ and floated on a 10 μL drop of staining solution (0.75% uranyl formate in $H_2O$) for 35 seconds. After removal of excess staining solution, samples were used for TEM analysis at 80 kV.

Thin section TEM analysis of fixed bacterial cells

For TEM analysis of fixed cells, 0.5 mL of early stationary phase bacterial culture was fixed by adding fixative (1:1 v/v, 1.25% formaldehyde, 2.5% glutaraldehyde, 0.03% picric acid in 0.1 M sodium cacodylate buffer, pH 7.4). The sample was then incubated at 25°C for 1 h and centrifuged for 3 min at 3000 rpm. The sample was then further incubated for 6-18 h at 4°C. Cells were subsequently washed three times in cacodylate buffer, 4 times with maleate buffer pH 5.15 followed by staining with 1% uranyl acetate for 30 min. The sample was dehydrated (15 min 70% ethanol, 15 min 90% ethanol, 2 x 15 min 100% ethanol) and exposed to propyleneoxide for 1 h. For infiltration, a mixture of Epon resin and proylenoxide (1:1) was incubated for 2 h at 25°C before moving it to an embedding mold filled with freshly mixed Epon. The sample was allowed to sink and subsequently moved to a polymerization oven (24 h, 60°C). Ultrathin sections (60-90 nm) were then cut at -120°C using a cryo-diamond knife (Reichert cryo-ultramicrotome) and transferred to formvar/carbon coated grids.

Cryo-electron microscopy (cryo-EM) data collection and processing

To prepare grids for cryo-EM imaging, 2.5 μL of purified cargo-loaded IMEF encapsulin at a concentration of 1.5 mg/mL was applied to glow-discharged Quantifoil holey carbon grids (1.2/1.3, 400 mesh), and blotted for 3 seconds with ~90% humidity before plunge-freezing in liquid ethane using a Cryoplunge 3 System (CP3, Gatan). Cryo-EM images were collected at Harvard Medical School on a Tecnai F20 electron microscope (FEI) operating at 200 kV and equipped with a K2 Summit direct electron detector (Gatan). Movies were collected at a nominal magnification of 29,000 with a calibrated pixel size of 0.64 Å. All movies were collected in super-resolution counting mode using UCSFImage4, with a total exposure time of 7.2 seconds and a frame time of 200 milliseconds. The details of EM data collection parameters are listed in Table S2.

Dose-fractionated super-resolution movies collected on the K2 detector were binned over 2 x 2 pixels, and subjected to motion correction using the program MotionCor2 (*29*). Dose-weighted sums from all frames were used for all subsequent image-processing steps except for defocus determination. The CTFFIND4 program (*30*) was used to determine the defocus values of the summed images from all movie frames without dose weighting. Semi-automated particle picking from 6x binned images was performed with SAMUEL and SamViewer (*31*). Selected particles were extracted from unbinned images with an initial box size of 512 pixels, and subsequently binned to a box size of 128 pixels with a pixel size of 5.12 Å for two rounds of 2D classification using RELION 3.0 (*32*). An initial 3D model was generated via SPIDER (*33*) 3D projection matching refinement (samrefine.py) using 2D class averages, starting from a sphere density similar in size and shape of the IMEF encapsulin. The selected particles after 2D classification were binned to a box size of 480 pixels (corresponding to a pixel size of 1.365 Å) and used for 3D refinement in RELION 3.0 with icosahedral symmetry ("I") imposed. A final round of 3D refinement was performed in RELION 3.0 after fitting individual particle defocus parameters and beam-tilt with "relion_ctf_refine". Post-processing was performed with "relion_postprocess" to apply a negative b-factor and correct the amplitude information in the final map. The overall resolutions were estimated based on the gold-standard criterion of Fourier shell correlation (FSC) = 0.143. Local resolution variations were estimated from two half data maps using ResMap (*34*).

Cryo-EM model building and refinement

An initial model of an IMEF encapsulin monomer was generated by homology modelling with the I-TASSER webserver (*35*) using the x-ray crystal structure of the T = 3 *Pyrococcus furiosus* encapsulin (PDB ID: 2E0Z) as a template. The monomer model was then fit into the 3D map in UCSF Chimera (*36*), and subsequently adjusted manually in COOT (*37*) prior to refinement in PHENIX (*38*) with phenix.real_space_refine. The refined monomer coordinates were copied and manually positioned to occupy the 4 monomer positions of the asymmetric unit (ASU), followed by manual adjustment of each monomer in COOT. Several rounds of real-space refinement and manual adjustment of the coordinates for four monomers in the ASU were performed in phenix.real_space_refine and COOT. During refinement of coordinates in the ASU no non-crystallographic symmetry restraints were utilized in order to avoid distortion of the E-loop in each monomer. The refined coordinates for the ASU were subsequently expanded using the symmetry matrices utilized by RELION 3.0 during 3D reconstruction to generate a model of the entire encapsulin cage containing 60 ASUs and 240 total IMEF encapsulin capsid protein polypeptide chains. Coordinates for the entire IMEF encapsulin cage were refined in phenix.real_space_refine with proper NCS restraints between corresponding chains in individual ASUs in order to resolve any inter-protomer clashes.

Differential scanning fluorimetry (DSF) to test thermal stability of proteins

DSF measurements were performed using a NanoTemper Tycho NT.6 instrument according to the manufacturer's instructions. Samples in Tris buffer (50 mM Tris, 150 mM NaCl, pH 8) at a concentration of 0.5 mg/mL were measured in triplicate and subjected to a temperature gradient from 35 to 95°C at 0.5°C per second. Data was analyzed using NT Melting Control software. Melting temperatures (Tm) were determined by automatic fitting of experimental data using a polynomial function, where the maximum slope (Tm) is indicated by the peak of its first derivative.

Crystallization and x-ray structure determination of the IMEF cargo protein

Initial crystallization conditions were determined using the Midas screen (*39*). Large single crystals were grown in sitting drop plates by the vapor diffusion method. Reservoir solutions contained 10% v/v Pentaerythritol ethoxylate (3/4 EO/OH) and 10% butanol. Crystals were cryo-protected in reservoir solution supplemented with 15% ethylene glycol and 20 mM glycolic acid pH 7.5. Diffraction data were collected at the European Synchrotron Radiation Facility (ESRF) Grenoble outstation at the ID-30b beamline at 100 K with a Pilatus3 6M pixel detector (DECTRIS, Switzerland). Data were indexed, processed, and scaled with the XDS package (*40*). The structure was solved by molecular replacement using an I-TASSER homology model and the program ACRIMBOLDO_LITE (*41*) incorporating PHASER (*42*) and SHELX (*43*) from the CCP4 suite (*44*). Model building and refinement was carried using COOT (*45*) and REFMAC5 (*46*),respectively.

Determination of electron-dense core diameters

To determine the size distribution of electron-dense cores resulting from IMEF mineralization under high iron conditions, TEM micrographs were analyzed using the open source image processing package Fiji based on ImageJ 1.52h. Micrographs were converted to 8-bit binary images, thresholded and processed using the particle analyzer plugin. The diameters reported are based on Fiji Feret diameter output values.

In vivo mineralization of electron-dense particles

Overnight cultures were used to inoculate 500 mL LB medium (1:50) supplemented with ampicillin and grown at 37°C to an $OD_{600}$ of 0.5. Expression was induced with 0.05 mM IPTG. Cultures were incubated at 30°C for 2 h. LB medium was removed and replaced with fresh modified LB (LB + 50 mM Hepes, 4 mM Trisodium citrate, pH 7) supplemented with freshly prepared ammonium iron(II) sulfate ($Fe(NH_4)_2(SO_4)_2$, final concentration: 4 mM; stock solution: 400 mM in 0.1 M HCl). The cultures were then incubated at 30°C for 18 h and used for either the purification of iron-loaded encapsulin compartments or thin section TEM analysis.

Iron-rich core characterization via energy-dispersive x-ray spectroscopy (EDS) and electron energy loss spectroscopy (EELS)

TEM and high angle angular dark field (HAADF) STEM imaging and analysis were performed on a JEOL ARM 200F operated at 80 kV. EDS spectra were collected using an EDAX Octane W 100mm2 detector, and spectra analyzed post-collection both via TEAM software and offline using the k-ratio method (thin film approximation). EELS mapping data of the Fe L edge was acquired using a Gatan Enfinium spectrometer with dispersion 0.25eV/ch using DualEELS mode with simultaneous zero loss spectrum collection. EELS data was processed using the Gatan EELS analysis plug-in. The processing steps involved a Gaussian fitting of the zero loss peak, integrating under the FeL edge up to 780 eV after applying a power law or first order log-polynomial (whichever fit the background better, as this depended on local carbon contamination levels) and correcting for the Fe cross section of 2664.9 barns, from which the average number of Fe per $nm^2$ was calculated per pixel of data. These pixels were summed over the area of each particle to estimate the total number of Fe atoms. Errors in this measurement were calculated from a statistical analysis of the data fitting combined with the expected error from Fe cross sectional extrapolation. Particle diameters were estimated using a histogram method to determine the edge onset of each particle, with the mean of multiple measurements from each particle used (and error determined by the standard deviation of these measurements).

Cultivation of *Geobacillus stearothermophilus* ATCC 7953

For normal growth of *G. stearothermophilus*, Meat Media (3 g meat extract, 5 g peptone, 1 L $H_2O$) was utilized. *G. stearothermophilus* was maintained on Meat Media agar plates (15 g agar/L). All growth was carried out a 55°C. For high iron growth experiments Meat Media was supplemented with 50 mM Hepes, 4 mM Trisodium citrate and 4 mM $Fe(NH_4)_2(SO_4)_2$ and the pH adjusted to 7 using HCl. Growth curves were recorded in high iron Meat Media (Fig. S17C) in 96-well plates (volume: 500 µL) using a Synergy H1 plate reader (BioTek) and inoculated (1:50) from a pre-culture grown for 24 h in standard Meat Media.

Peroxidase assays

Peroxidase activity of free IMEF cargo and cargo-loaded IMEF encapsulin was assayed by measuring the oxidation of *ortho*-phenylenediamine (OP) by hydrogen peroxide (*47*). OP dilutions from 10 to 80 mM were prepared from a stock solution (92.5 mM in 50 mM Tris, pH 8) using Tris buffer (pH 8). 96-well plates were used to carry out the assays in triplicate. Each well contained 100 µL of OP dilution and 0.5 µM of IMEF cargo protein (protein concentrations were determined via Bradford assay (Pierce Coomassie, ThermoFisher) following the manufacturer's instructions). To start the assays, 2 μL of 30% hydrogen peroxide solution was added. After 15 min of incubation in the dark, assays were stopped by the addition of 100 μL of 0.5 M $H_2SO_4$. Then, absorbance at 490 nm was determined using a Synergy H1 plate reader.

Ferroxidase assays

Protein solutions in Tris buffer (50 mM Tris, 150 mM NaCl, pH 8) and $Fe(NH_4)_2(SO_4)_2$ stock solutions in 0.1 M HCl were made anaerobic by incubation in a Vinyl Anaerobic Chamber (Coy Lab Products) for 24 h. All solutions were exposed to the anaerobic atmosphere inside the chamber and protein solutions were kept on ice. IMEF cargo protein was used at a final concentration of 50 μM while cargo-loaded encapsulin concentrations were used that would correspond to 5 μM IMEF cargo (higher concentrations led to rapid protein precipitation upon iron addition). Final iron(II) concentrations ranged from 10 to 100 μM. Ferroxidase activity was initiated by combining appropriate dilutions of protein and iron solution to a final volume of 250 μL in a quartz cuvette in the air, directly after removing solutions from the anaerobic chamber. Ferroxidase activity was immediately measured by monitoring $Fe^{3+}$ formation at a wavelength of 315 nm in a Nanodrop 2000c for 25 min.

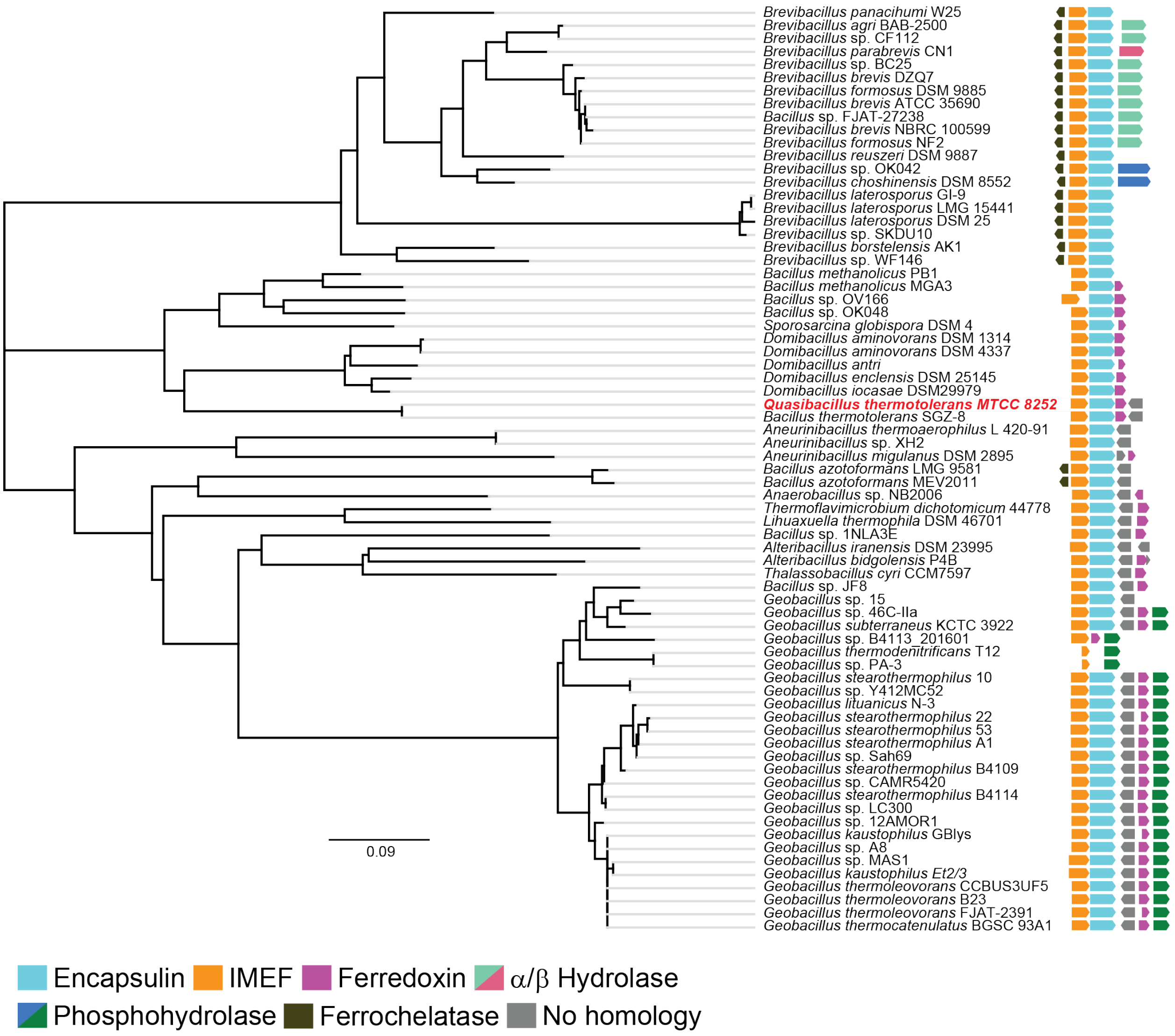


**Fig. S1. The 71 IMEF operons identified in Firmicutes.** The *Qs* operon is highlighted in red. The nearest-neighbor phylogenetic tree is based on a ClustalO (*48*) alignment of IMEF cargo proteins. IMEF cargo accessions are shown in Table S1. Evolutionary distances were estimated as the number of amino acid substitutions per site. The scale bar represents 0.09 expected amino acid residue substitutions per site.

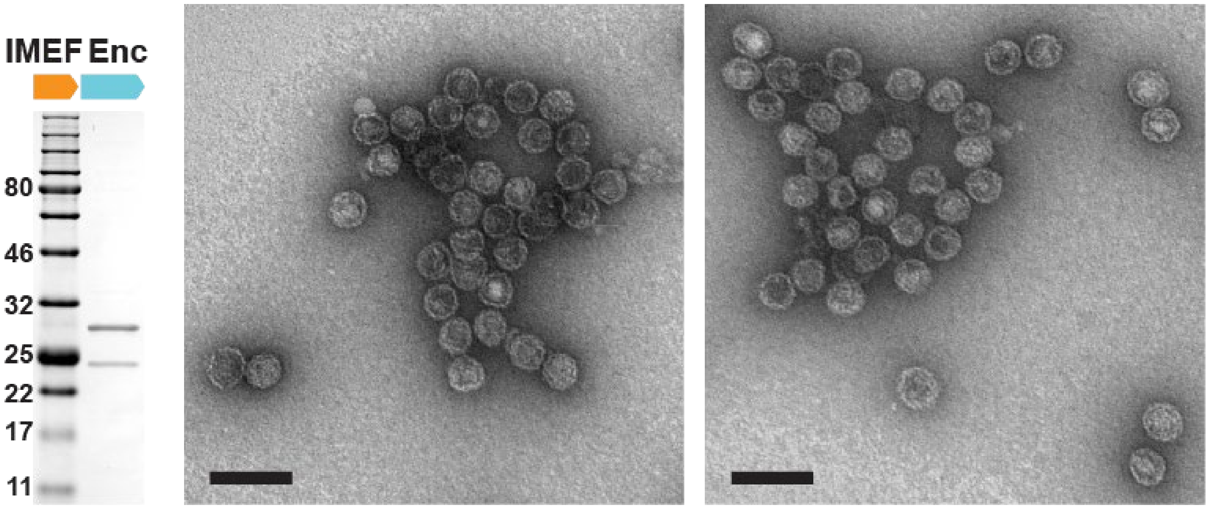


**Fig. S2. Recombinantly produced and purified cargo-loaded IMEF encapsulins.** Shown are negative stain (uranyl formate) TEM micrographs of purified compartments (right) and a representative SDS-PAGE gel (left). The SDS-PAGE gel shows that the IMEF cargo protein (22.6 kDa) co-purifies with the encapsulin capsid protein (32.2 kDa). The SDS-PAGE gel is the same as shown for comparison in Fig. 3E of the main text. Scale bars in micrographs correspond to 100 nm.

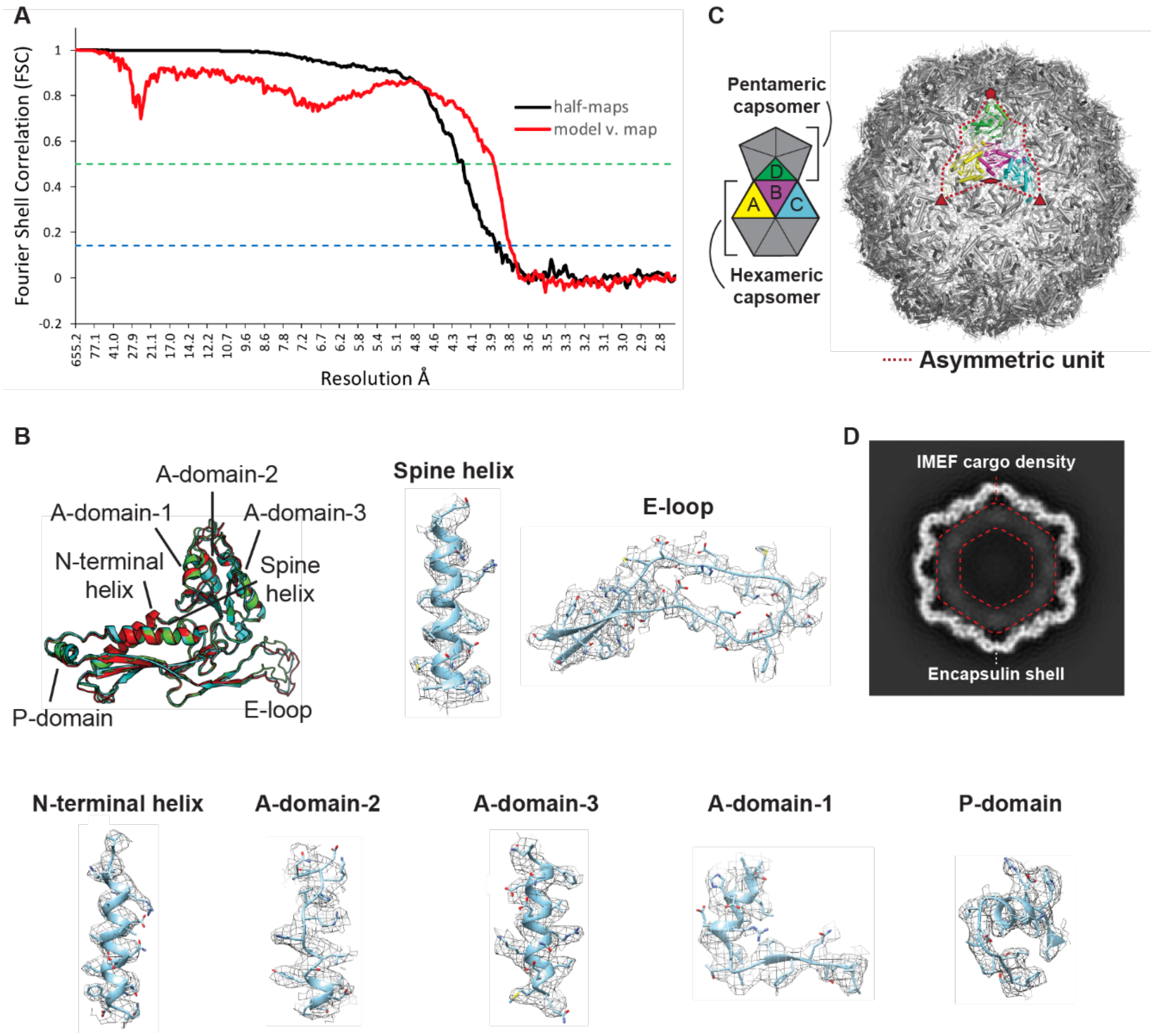


**Fig. S3. Cryo-EM data analysis and evaluation.** (**A**) FSC curves calculated from independent half-maps (black line) and a comparison of the atomic model of the IMEF encapsulin versus the experimental map (red line). The green dashed line represents an FSC cutoff of 0.5 and the blue dashed line represents an FSC cutoff of 0.143. (**B**) Annotated monomer alignment of the IMEF capsid protein monomers with local cryo-EM maps. The different structural elements of the T = 4 encapsulin monomer are shown. Representative density maps (mesh) and atomic models (model B of the asymmetric unit) illustrating side chain features and overall fit of the atomic model with the cryo-EM map are shown. (**C**) T = 4 IMEF encapsulin capsid with highlighted asymmetric unit (right) and annotation (left) defining the 4 capsid monomers of the asymmetric unit and the hexameric and pentameric capsomers. (**D**) Central slice of cargo-loaded IMEF encapsulin capsid. Less defined lower resolution density can clearly be seen in the interior (red dotted lines).

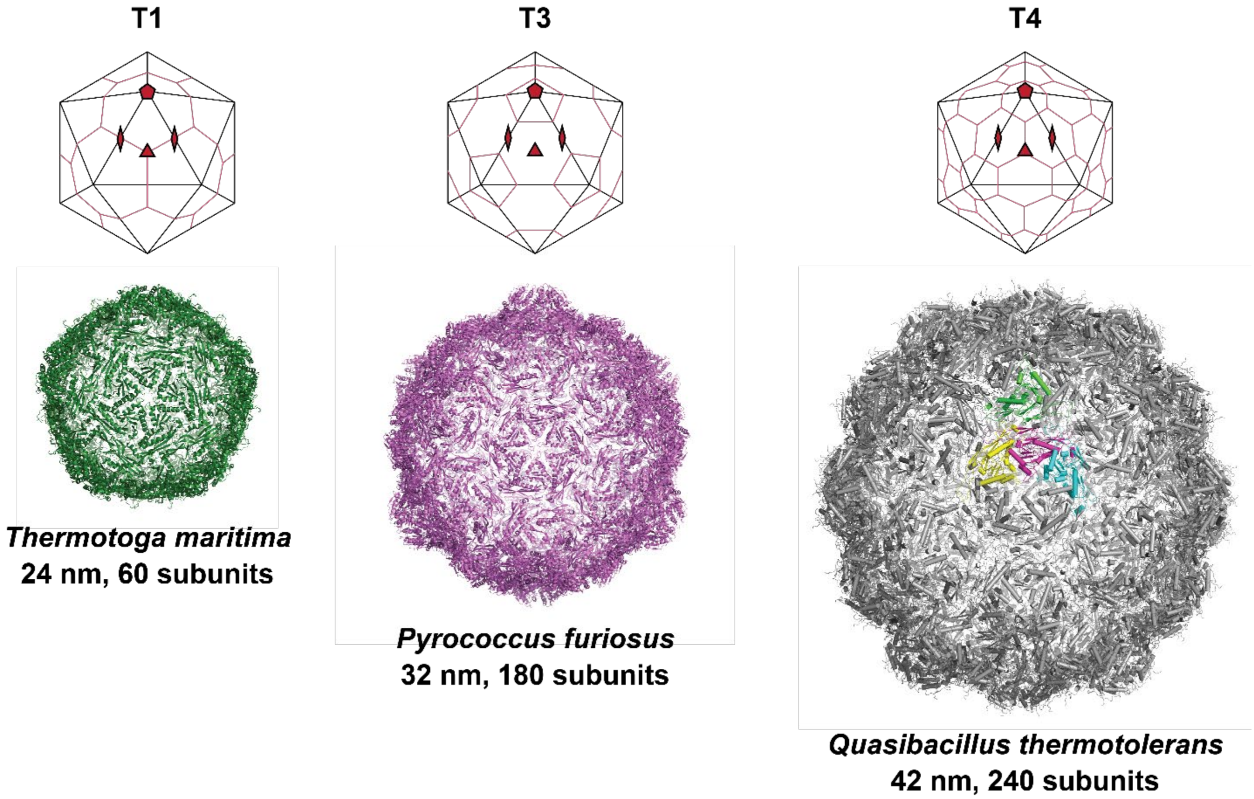


**Fig. S4. Structural and topological comparison of different encapsulins.** Top: Drawings illustrating the icosahedral symmetry and tiling of capsids with different triangulation numbers. Bottom: Capsid models of T = 1, T = 3 and the newly discovered T = 4 encapsulin shells including maximum external diameter and subunit number.

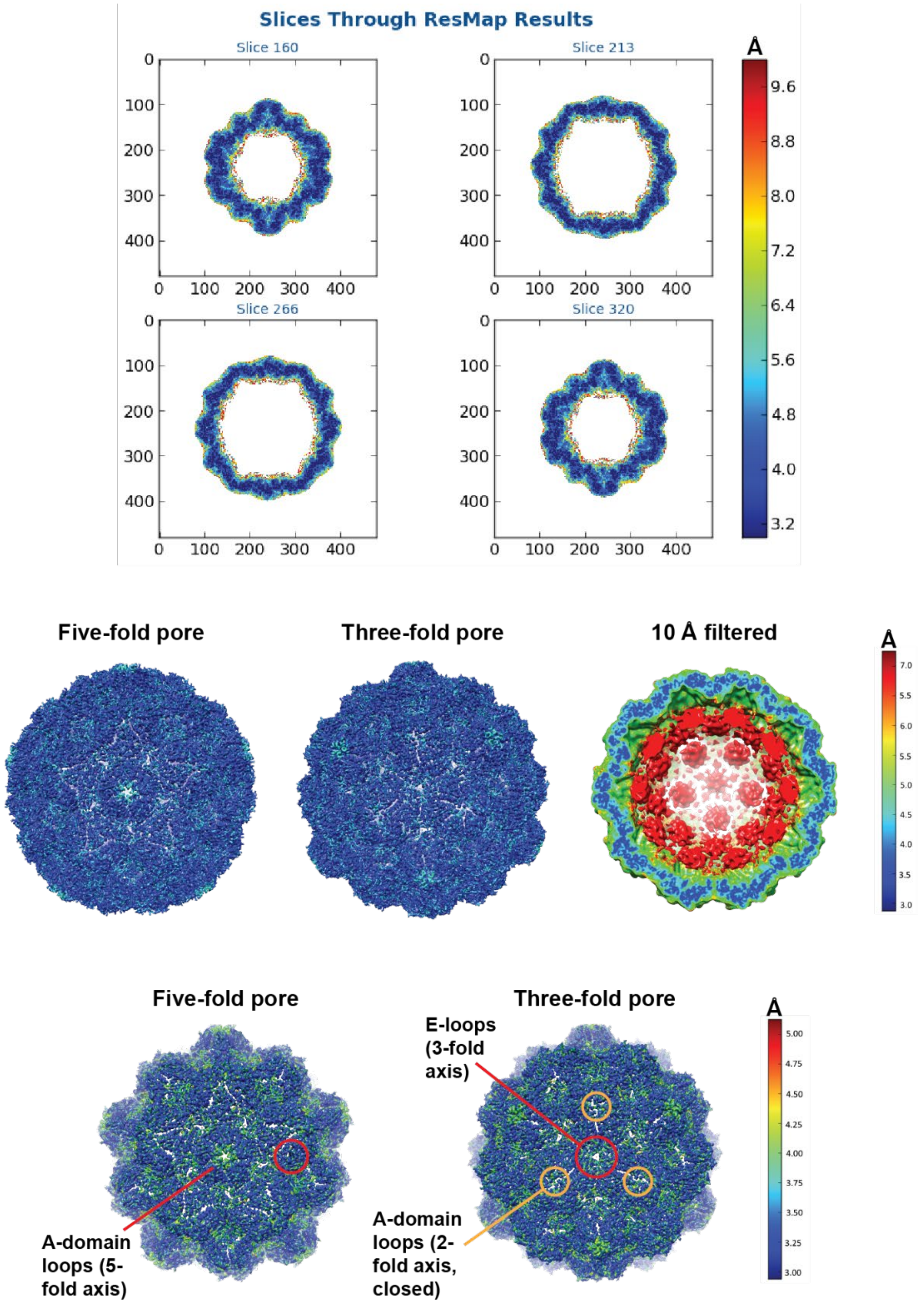


**Fig. S5. Local resolution maps of the cargo-loaded T = 4 IMEF encapsulin.** Top: central slices through the T = 4 shell. Middle: Comparison of shell and internal cargo resolution. Bottom: Focus on shell resolution. The lowest resolution parts of the T = 4 shell are highlighted and include 5- and 3-fold pore residues corresponding to A-domain loops and E-loops, respectively, and A-domain loops forming the closed ‘pore’ at the 2-fold symmetry axis.

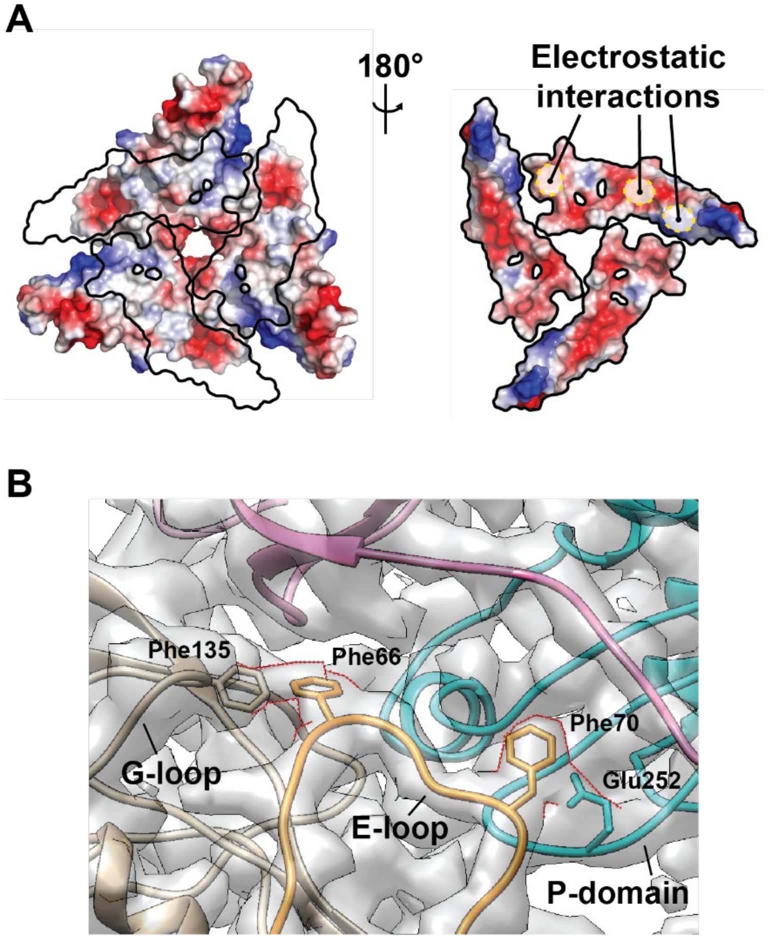


**Fig. S6. Subunit interactions at the 3-fold pores.** (**A**) Electrostatic surface models of P-domains and E-loops (outlines on the left and surface on the right) highlighting complementary electrostatic interactions around 3-fold pores. (**B**) Specific interactions observed at the interface of 3 capsid monomers. Strong cryo-EM density observed connecting subunits are outlined with red dotted lines. This density suggests aromatic interactions for Phe135 and Phe66 and potential anion-π interactions for Phe70 and Glu252. The interactions between the G-loop, E-loop and P-domain of 3 different subunits are likely one of the factors responsible for the observed thermal stability of this system. It is of note that the proposed Phe70-Glu252 interaction in the T = 4 IMEF encapsulin is located at the same location as the isopeptide bond observed in the HK97 bacteriophage capsid.

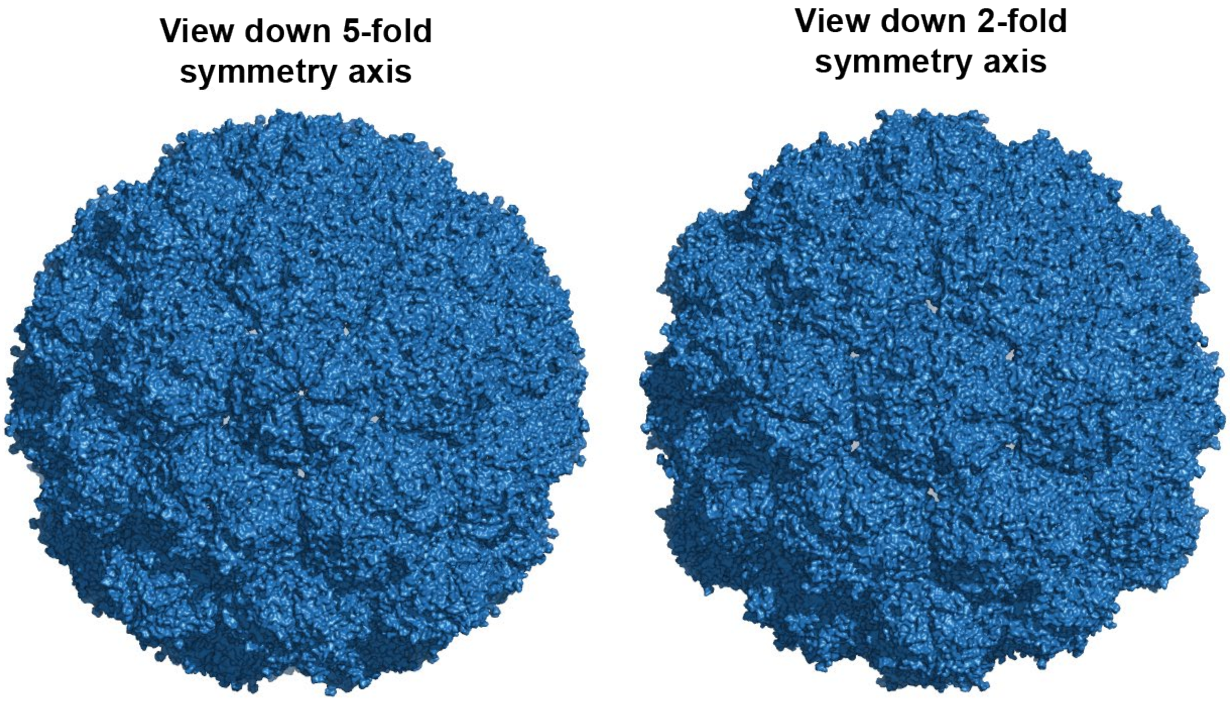


**Fig. S7. Surface views of the T = 4 IMEF encapsulin shell.** Left: View down the 5-fold symmetry axis highlighting the 5-fold and pseudo 3-fold pores. Right: View down the 2-fold symmetry axis highlighting the 3-fold pores and the closed 2-fold 'pore'. This surface representation indicates tight packing of monomers resulting in the 5- and (pseudo) 3-fold pores being the only conduits to the internal space created by the encapsulin shell. Pseudo 3-fold pores are defined as pores at the interface of 2 hexameric and one pentameric capsomer that in contrast to actual 3-fold pores (the interface of 3 hexameric capsomers) do not coincide with an icosahedral 3-fold symmetry axis.

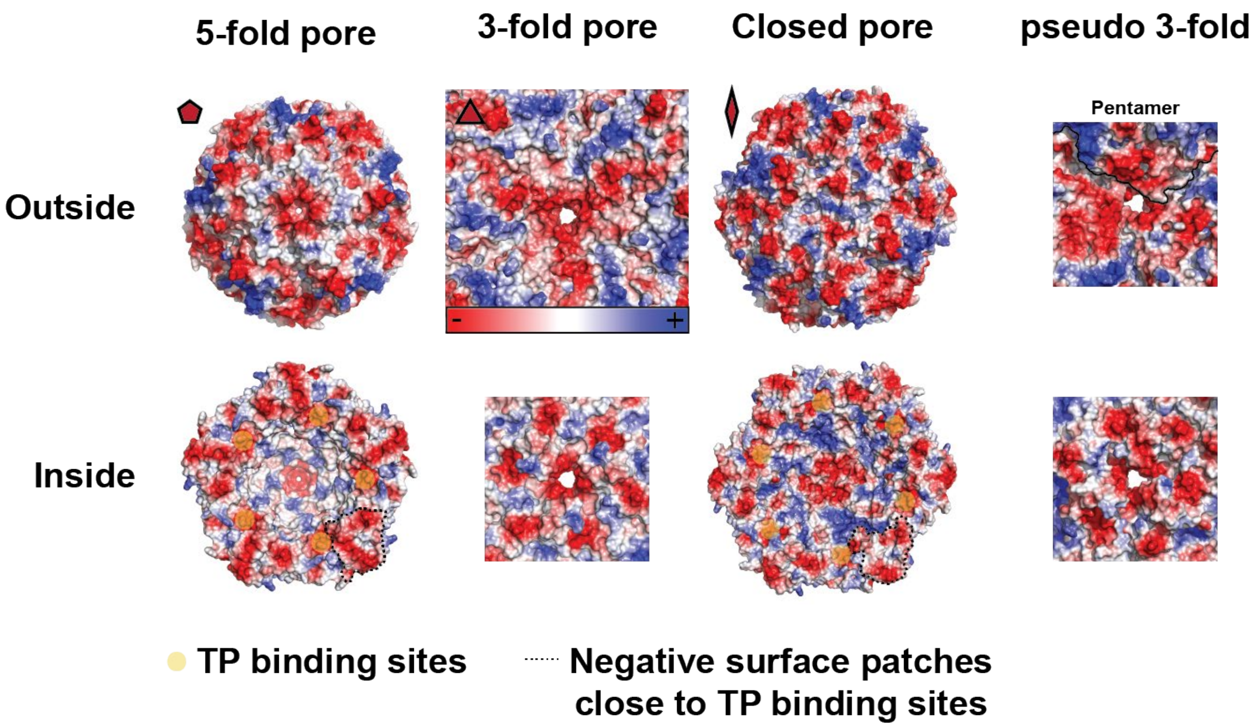


**Fig. S8. Overview of T = 4 IMEF encapsulin pores.** Electrostatic surface representations of both the inside and outside views are shown. 5- and (pseudo) 3-fold pores are clearly negatively charged on both sides and all the way through the pore itself. The potential pore at the 2-fold symmetry axis is positively charged (outside) right at the pore entrance due to the presence of two asparagine residues closing off the pore while the inside of the 2-fold 'pore' is strongly negatively charged similar to the other pores. TP binding sites around the 5- and 2-fold symmetry axes are indicated with yellow circles and large negatively charged surface patches close to TP binding sites are outlined by black dotted lines. These patches might be involved in increasing cargo affinity for the interior encapsulin shell close to the binding site due to ionic interactions with positively charged residues of the long IMEF cargo linker.

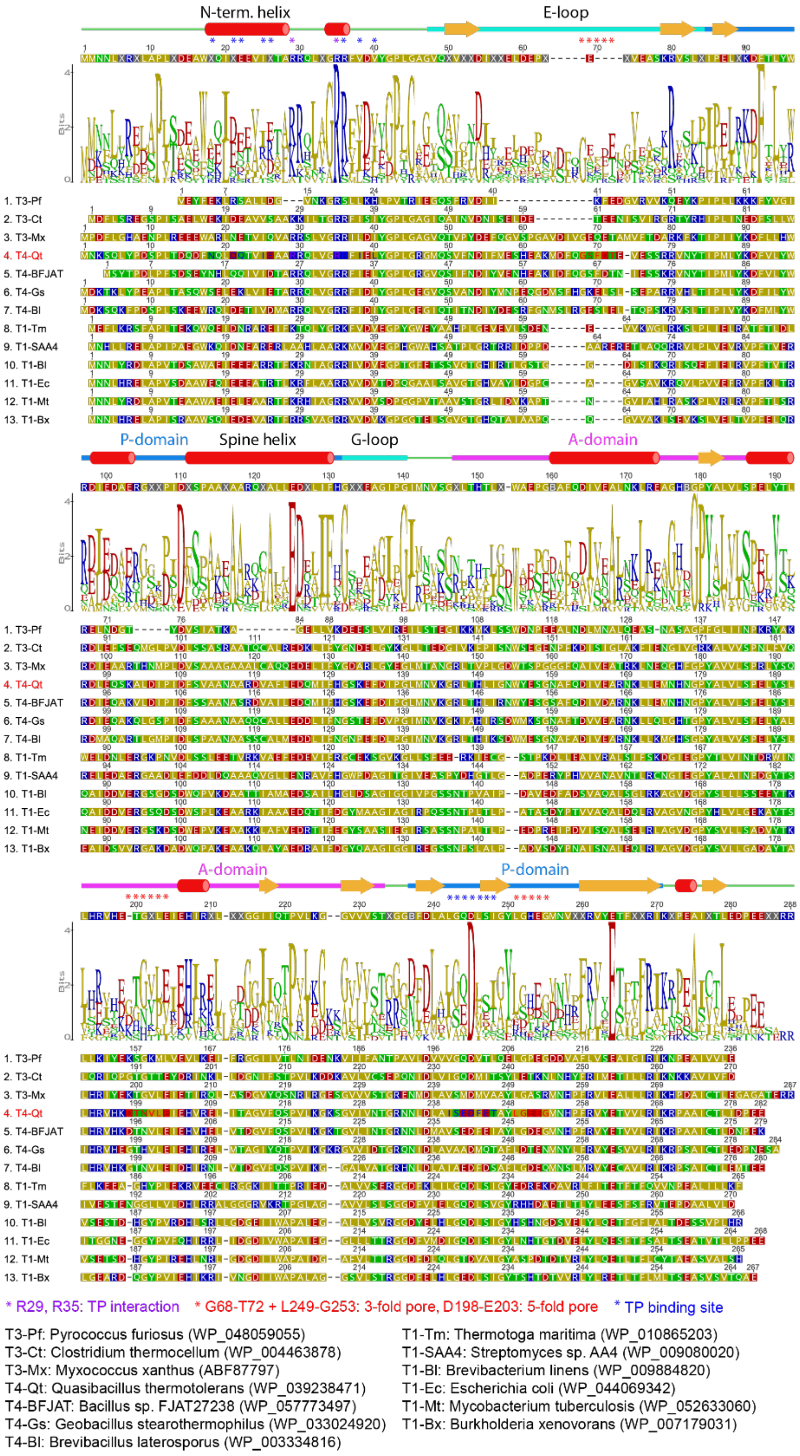


**Fig. S9. Sequence alignment of representative T = 1, T = 3 and T = 4 encapsulin capsid proteins.** Annotated secondary structural elements, the consensus sequence and a sequence logo are shown above the sequences. Organisms and protein sequences used are shown below the alignment. Residues important for TP interaction (purple), formation of the 3- and 5-fold pores (red) and in forming the overall TP binding site (blue) are indicated with asterisks.

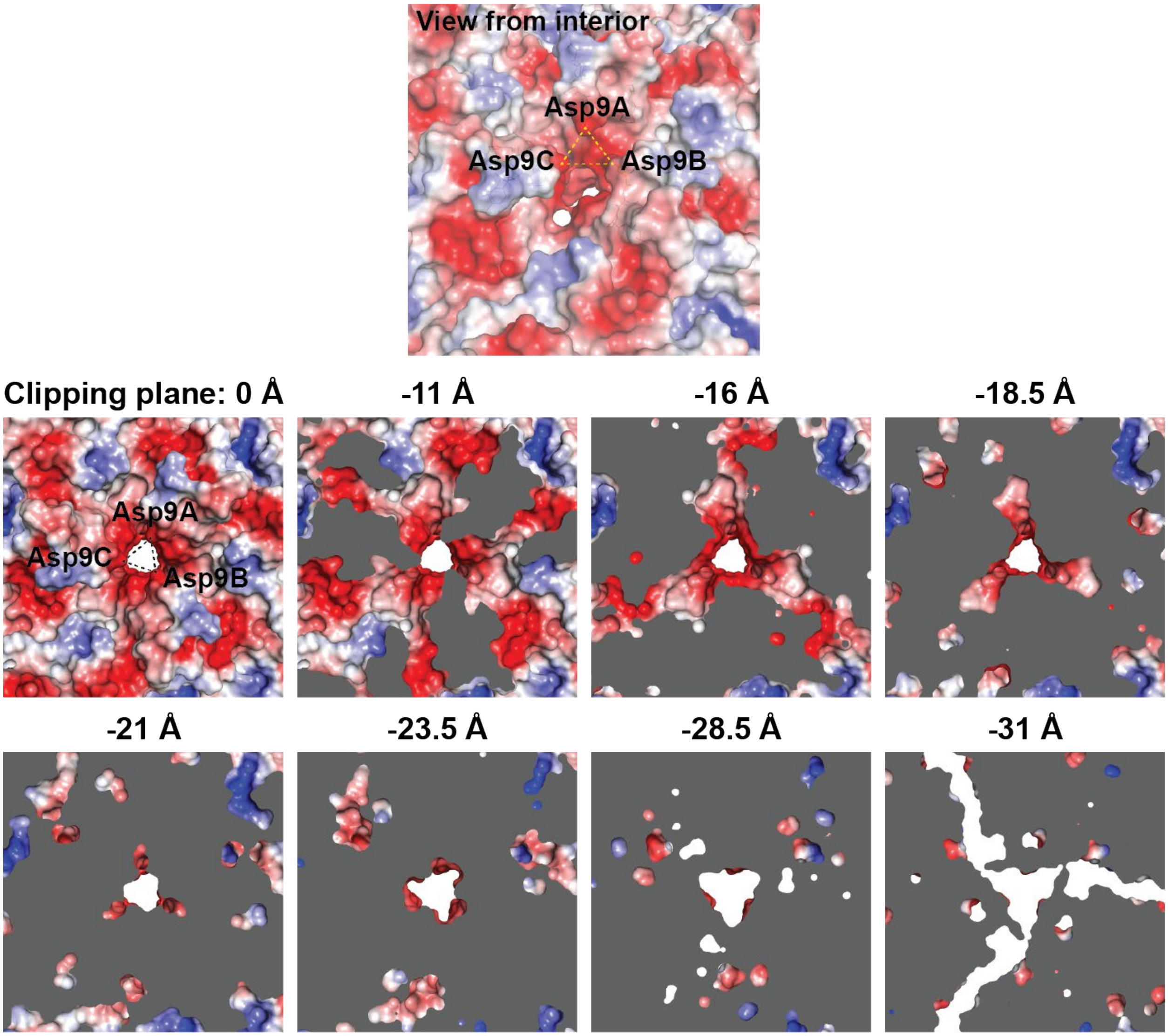


**Fig. S10. Detailed view of the large 3-fold pore.** Top: View from the inside highlighting negatively charged aspartate residues placed directly above the pore entrance. Bottom: Sequence of slices perpendicular to the 3-fold pore highlighting its overall shape and narrowest point which is still the largest pore observed for any encapsulin (7.2 Å, see Fig. 2 in the main text).

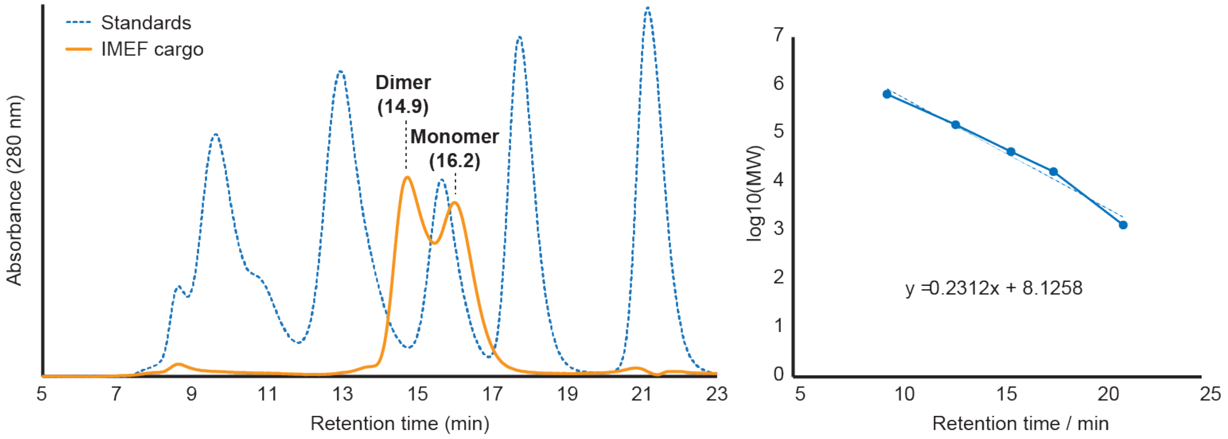


**Fig. S11. Analytical gel filtration of the IMEF cargo protein.** Left: Chromatogram indicating that His-tagged IMEF exists as a mixture of dimer and monomer in solution (in the absence of encapsulin). Right: Calibration curve used to estimate molecular weights and oligomerization state.

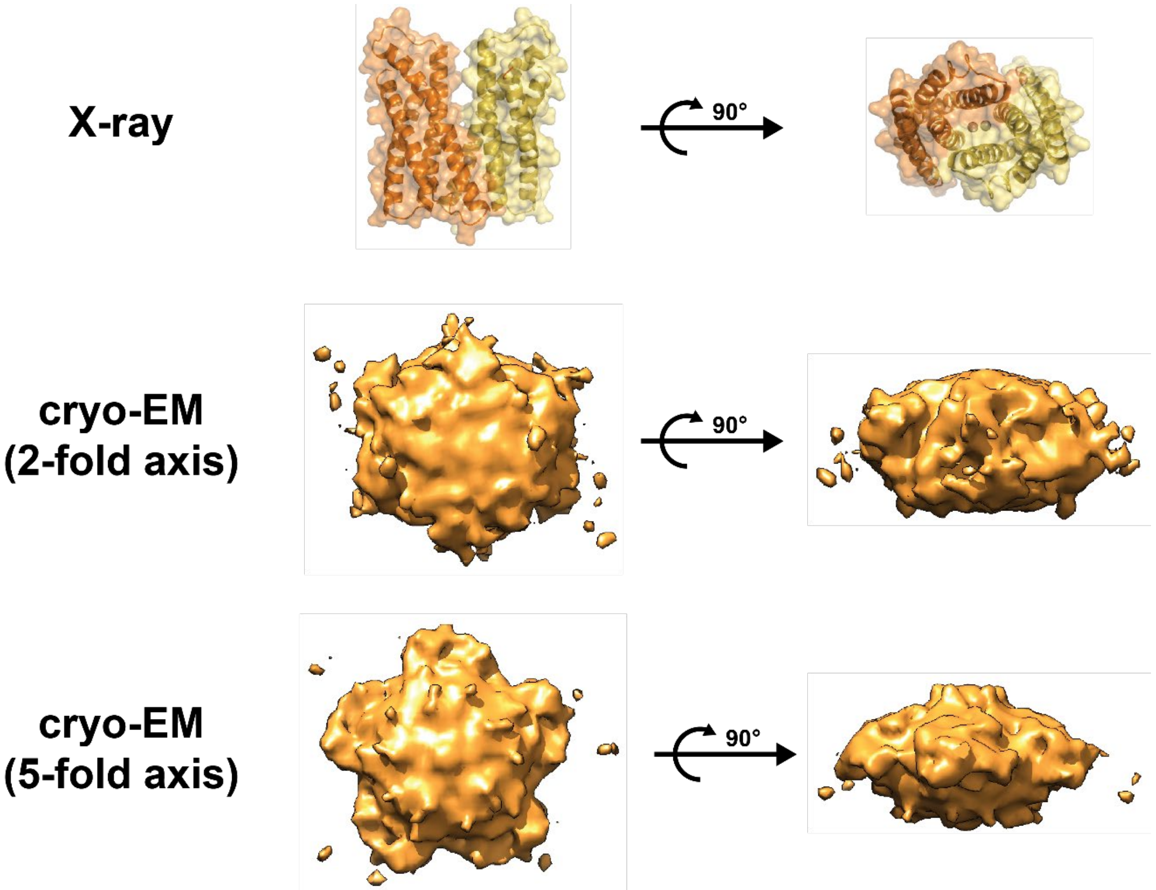


**Fig. S12. X-ray and cryo-EM IMEF cargo structure comparison.** Top: X-ray structure of the dimeric IMEF cargo. Middle: Cargo density observed above hexameric capsomers (2-fold symmetry axis). Bottom: Cargo density observed above pentameric capsomers (5-fold symmetry axis). The symmetry of cryo-EM cargo densities indicates averaging during cryo-EM reconstruction. The sizes of the densities observed in combination with x-ray and gel filtration experiments suggests that the IMEF cargo is present in a dimeric form when encapsulated. Due to steric hinderance it is very unlikely that on average more than one dimer per hexameric or pentameric capsomer is present in a fully loaded IMEF encapsulin. This means that when fully loaded 42 IMEF dimers are present per T = 4 IMEF encapsulin.

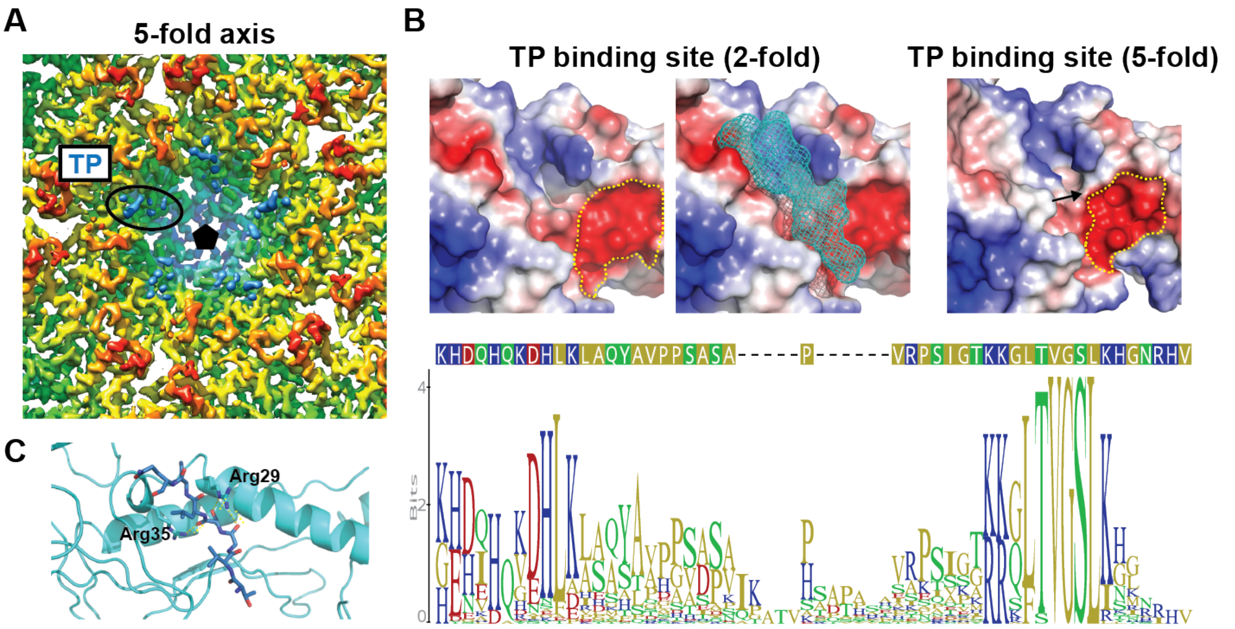


**Fig. S13. Analysis of TP binding sites.** (**A**) TP densities observed around the 5-fold symmetry axis. The densities are weaker than densities observed around 2-fold symmetry axes (Fig. 3F). (**B**) Top: 2- and 5-fold TP binding sites. A mesh model of the modelled TP is shown in cyan. Yellow dotted lines highlight large negative surface patches. Conformational changes lead to a less pronounced surface groove for the 5-fold binding site (black arrow). Overall the 2- and 5-fold binding sites are different due to the different conformations of capsid monomers when present in hexameric vs. pentameric capsomers, explaining the different binding affinities and observed differing TP density strengths. Bottom: Sequence logo and consensus sequence of the flexible linker and TP of all identified IMEF cargo proteins indicating the presence of many positively charged residues at both ends of the linker connecting IMEF and TP. (**C**) TP binding site (2-fold) highlighting key ionic interactions. The shell monomer is shown as ribbons, the TP is shown in stick representation.

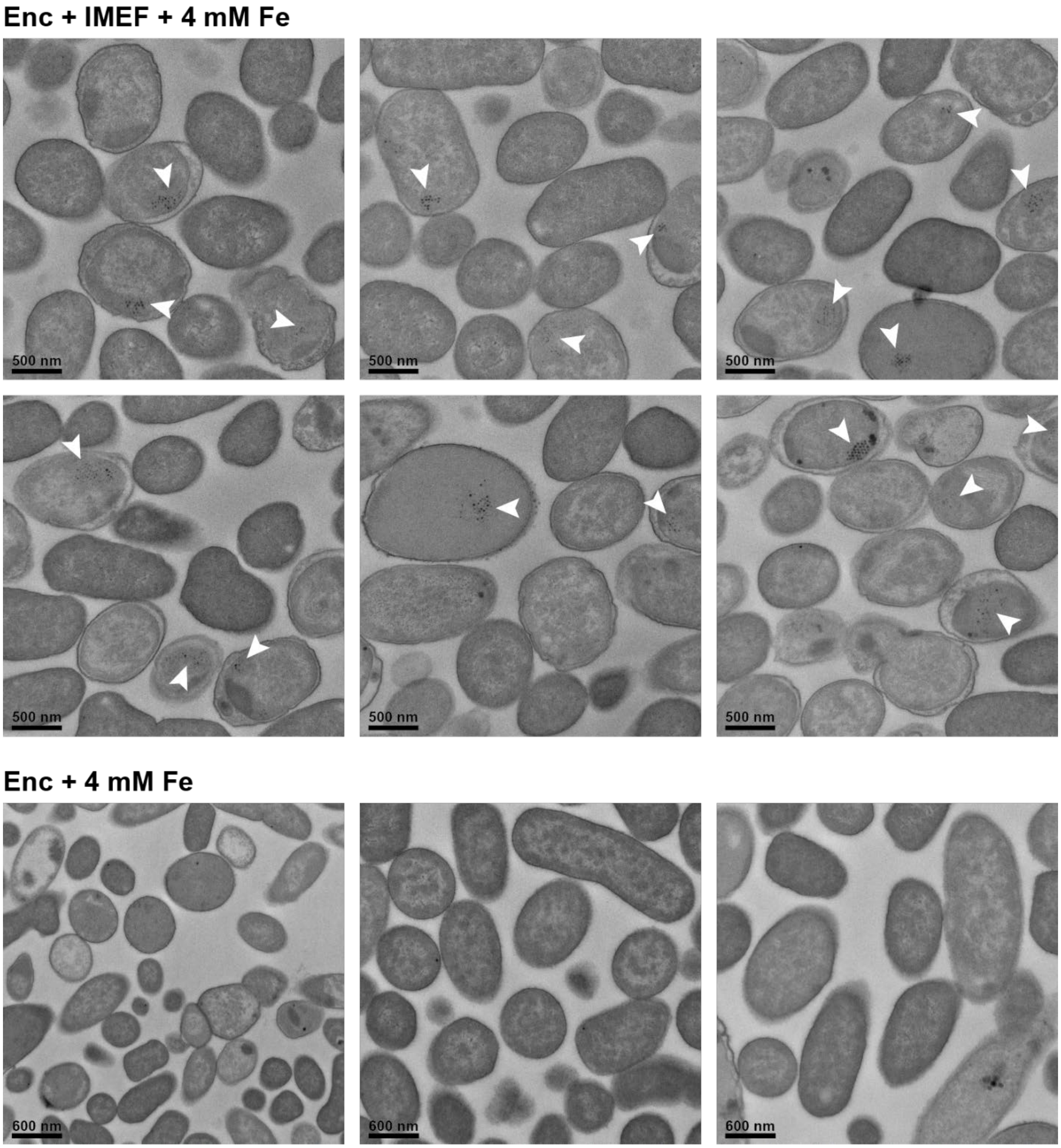


**Fig. S14. Negative stain thin section TEM micrographs of *E. coli*.** Top: Representative fields of view for *E. coli* expressing the core IMEF operon (IMEF cargo + capsid protein) under high iron conditions (4 mM $Fe(NH_4)_2(SO_4)_2$). Clusters of electron-dense particles are highlighted with white arrows. Bottom: E. coli cells expressing only the IMEF encapsulin capsid protein without the IMEF cargo. No clusters of electron-dense particles were observed.

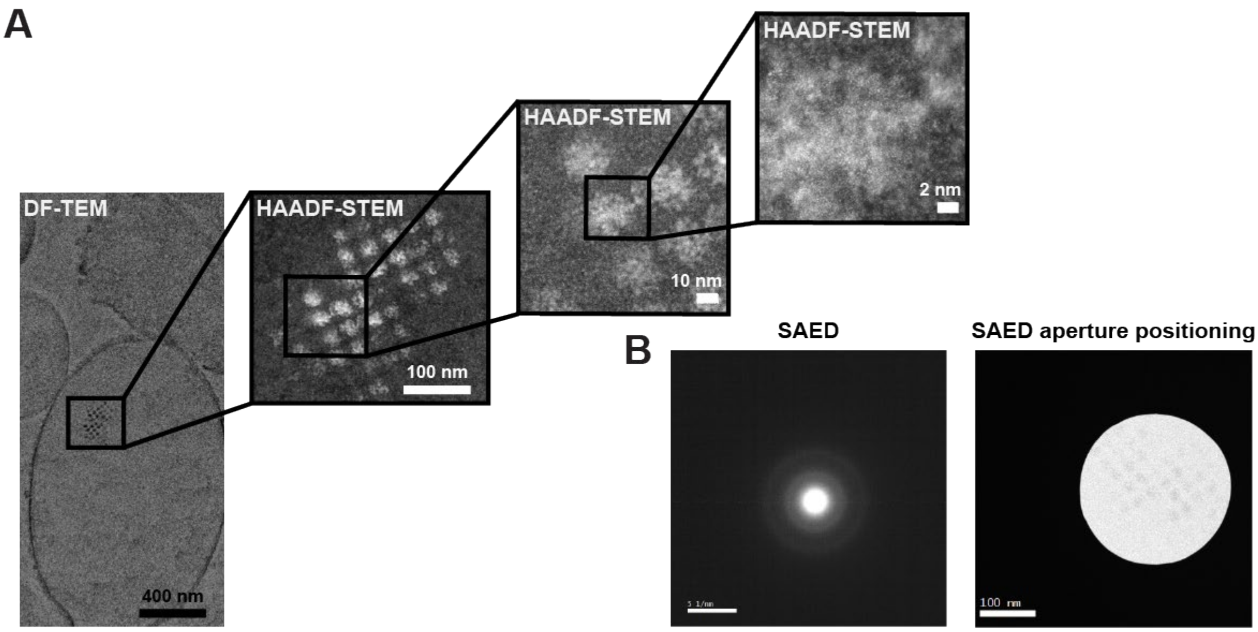


**Fig. S15. Mineralized iron-rich cores are amorphous.** (**A**) HAADF-STEM images at high resolution show irregular non-crystalline material. (**B**)Selected area electron diffraction (SAED) of electron-dense particles was carried out on thin sections of *E. coli* expressing the core IMEF operon targeting clusters of electron-dense particles. No diffraction spots and thus no crystallinity could be observed meaning that the electron-dense material deposited inside IMEF encapsulin shells is amorphous (supported by the presence of faint concentric rings indicative of an amorphous phase). HAADF-STEM: high angle angular dark field scanning TEM. DF-TEM: dark field TEM.

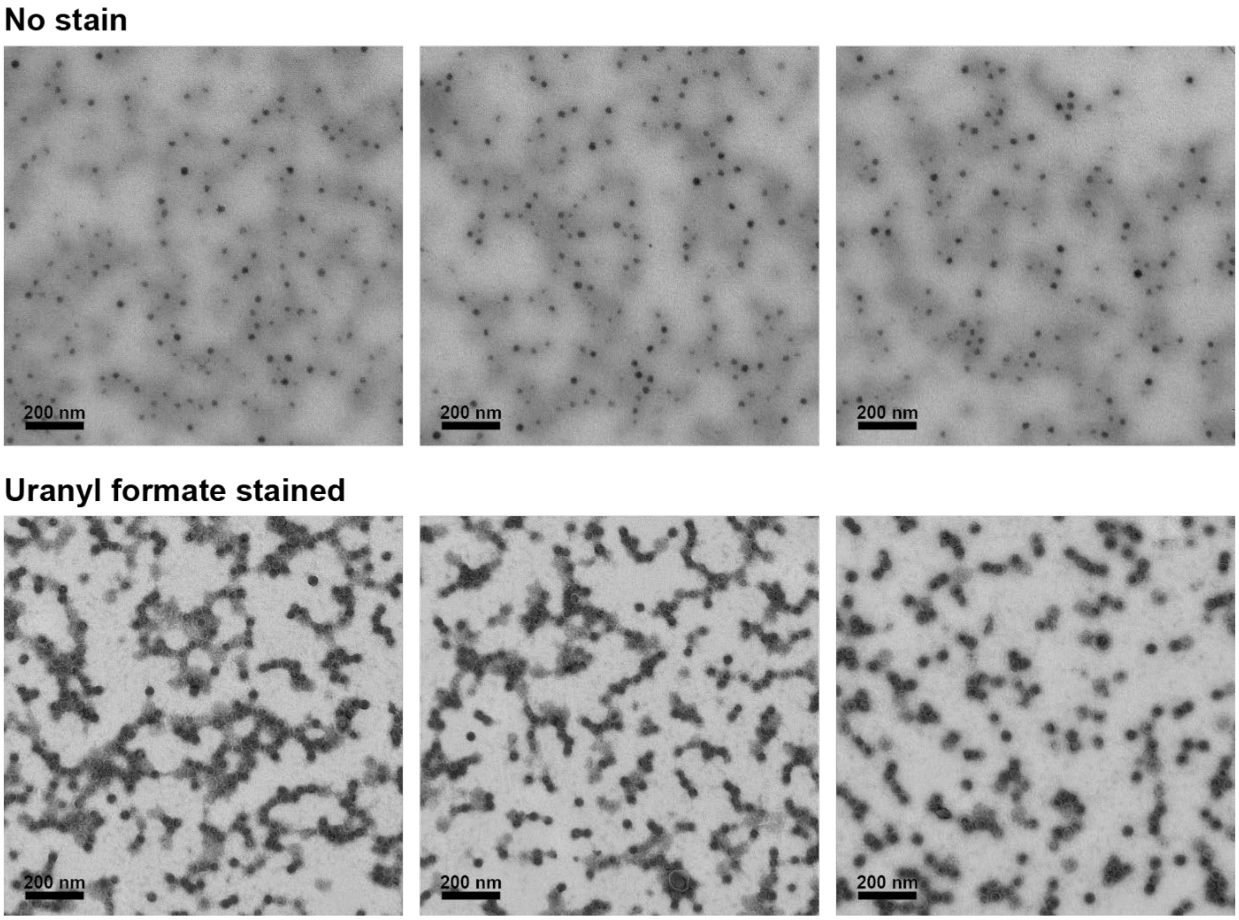


**Fig. S16. TEM micrographs of purified IMEF encapsulins produced under high iron conditions.** Top: Representative fields of unstained particles. Electron-dense cores are clearly visible even without stain. Bottom: Uranyl formate stained representative fields of the same particles. Images as shown on top were used to determine the core size distribution using ImageJ.

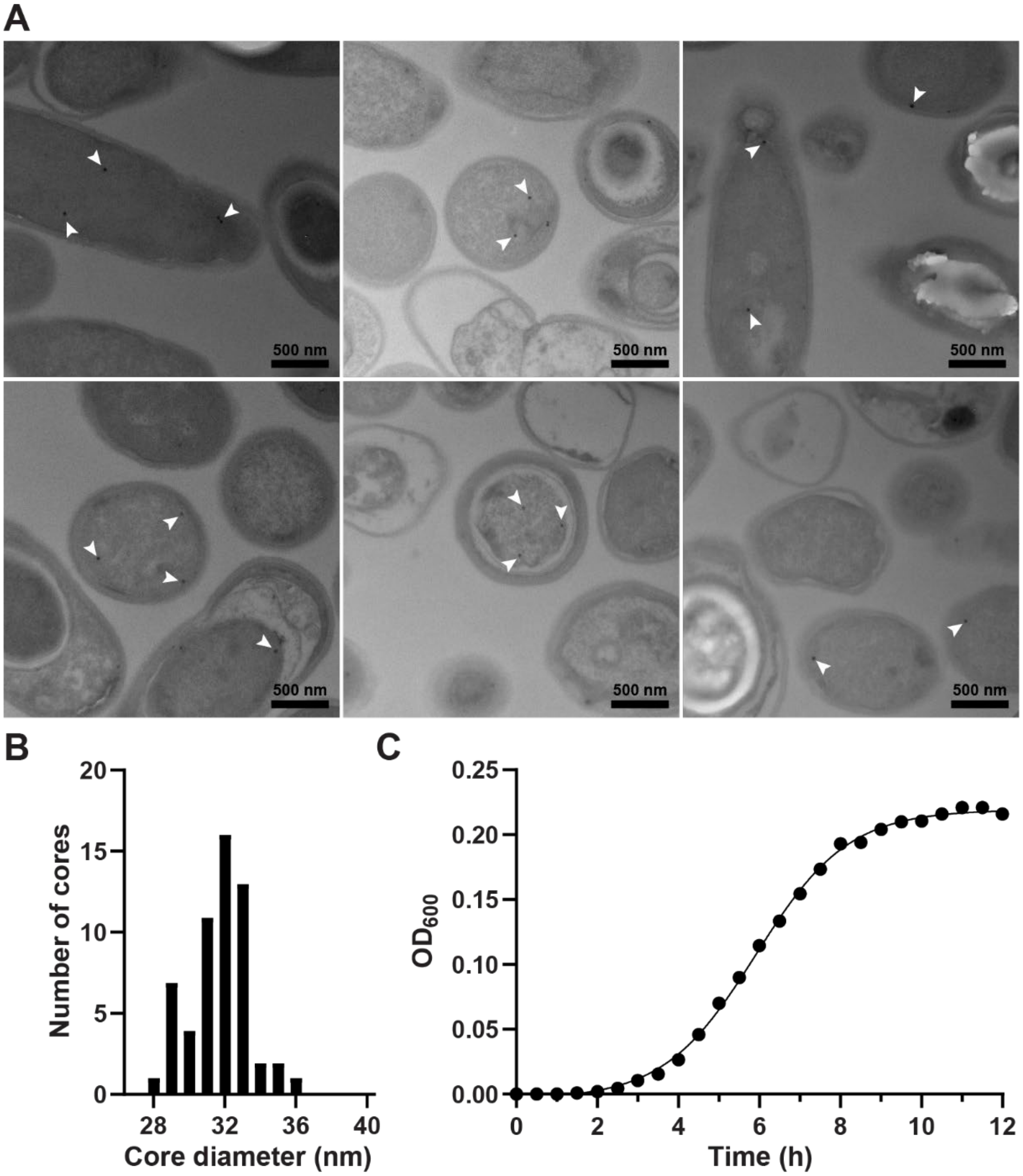


**Fig. S17. Iron-core mineralization in *Geobacillus stearothermophilus* ATCC 7953.** (**A**) *G. stearothermophilus* ATCC 7953 was grown in high iron medium (4 mM) and samples prepared in early stationary phase (after 9 h). Representative fields of cells are shown with electron-dense cores highlighted by white arrows. Substantially fewer cores were observed in this strain that natively encodes the IMEF operon compared with recombinant *E. coli* heterologously expressing the IMEF operon. Consequently, no clustering of cores was observed. These thin section micrographs were not suitable for more detailed core analyses like EDS and EELS due to rapid carbon build-up and very high background. Thus, materials characterization of cores was done on purified iron-loaded particles that could be isolated in high quantity from recombinant *E. coli* resulting in less carbon build-up and better signal-to-noise. (**B**) Size distribution of native electron-dense cores formed in *G. stearothermophilus*. (**C**) Representative growth curve of *G. stearothermophilus*.

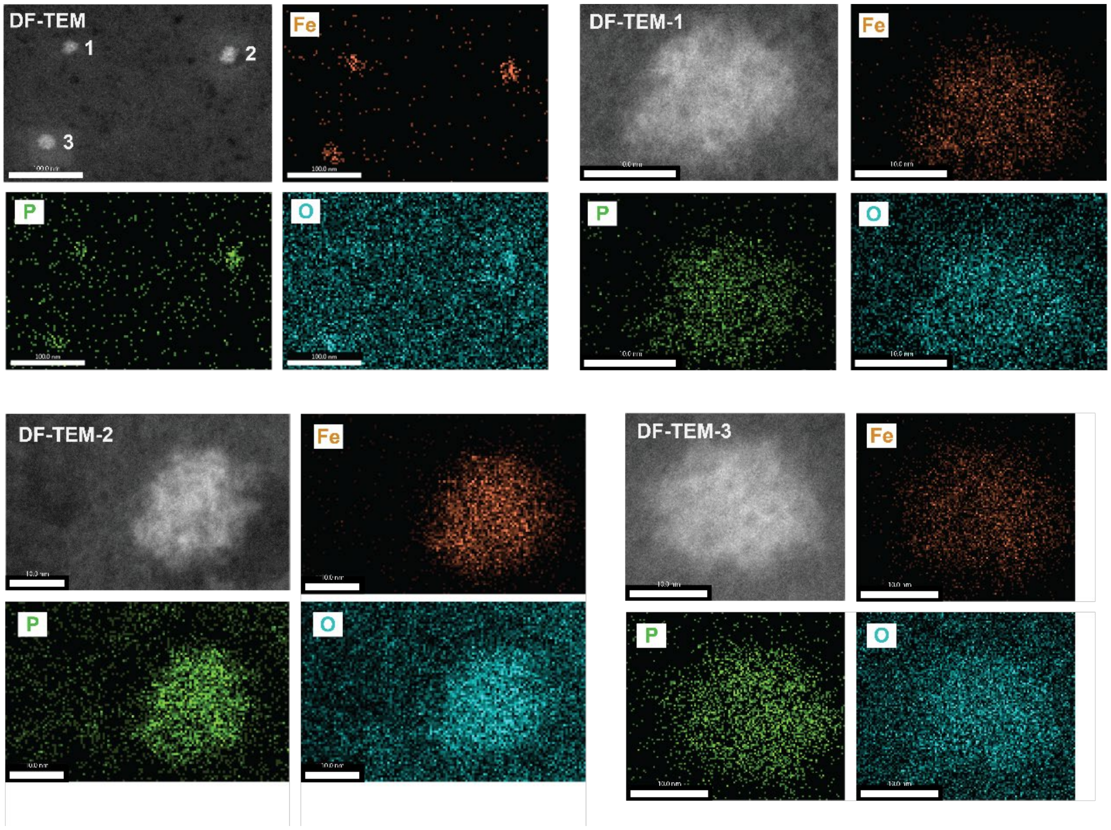


**Fig. S18. EDS analysis of purified iron-loaded encapsulin particles.** Three representative particles are shown with individual elemental EDS maps for Fe, P and O shown in orange, green and cyan, respectively.

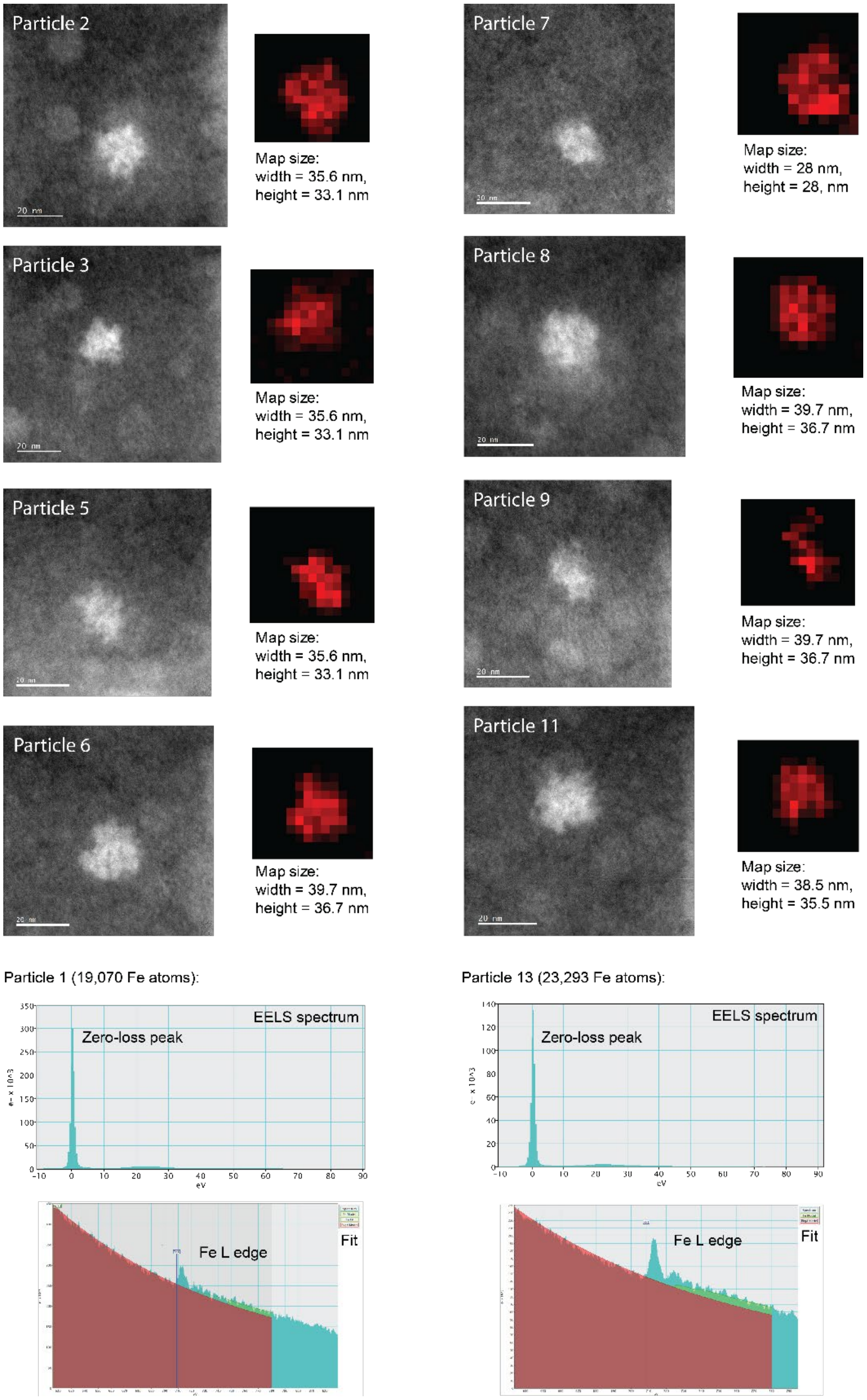


**Fig. S19. EELS analysis of purified iron-loaded encapsulin particles.** Top: representative particles (HAADF-STEM images) and corresponding EELS maps are shown. Bottom: EELS spectra and fit for particle 1 and particle 13 (shown in Fig. 4E and F in the main text, see also: Table S5).

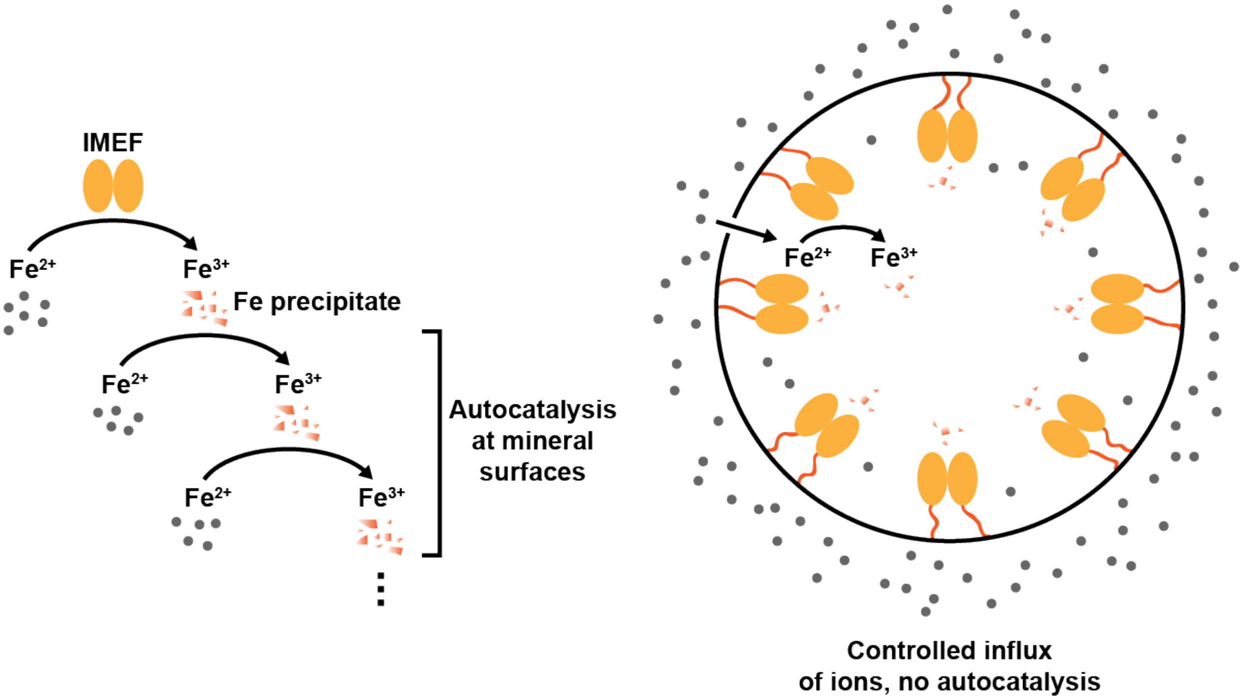


**Fig. S20. Model explaining the observed ferroxidase activities of free IMEF cargo (left) and the cargo-loaded IMEF encapsulin (right).** Without a protein shell creating a barrier between the compartment interior and outside, autocatalytic oxidation of ferrous iron on enzymatically formed ferric iron precipitates is observed (left) leading to a sigmoidal time-course curve as shown in Fig. 4G in the main text. However, in the presence of a protein shell, a characteristic hyperbolic enzyme catalysis curve is observed (Fig. 4H). This suggests that the encapsulin shell strictly controls the influx of iron to the compartment interior and thus the internal concentration of iron substrate available to encapsulated IMEF cargo proteins. The overall effect of this arrangement is that mineralization inside the IMEF encapsulin is controlled and autocatalytic runaway iron precipitation prevented. This is likely of key importance for the functioning of this novel iron storage system in the bacterial cytoplasm resulting in an iron storage system able to safely store essential but also toxic iron in a soluble and bioavailable form.

**Table S1. Ferritin-like proteins (Flps) identified in IMEF operon-containing Firmicutes.** All identified ferritins (Ftn), bacterioferritins (Btf) and DNA-binding proteins from starved cells (Dps) found in IMEF operon strains. IMEF and encapsulin capsid protein IDs are shown as well. 97% of IMEF operon-containing strains do not encode Ftn, 93% do not encode Bfr and 92% do not encode either. However, 93% of IMEF operon-encoding strains encode Dps systems. This likely indicates that the encapsulin based IMEF system represents the major iron storage system in 92% of the listed strains. It also indicates that IMEF systems do likely not function as unusual Dps system given that the vast majority of strains encode standard Dps system. Blast searches were carried out using the NCBI Blastp server with the following sequences as queries: Ftn: OTY20392, Bfr: EEK74551, Dps: WP_039234032, IMEF: WP_039238473, Encapsulin: WP_039238471.

| Organism | Ftn | Bfr | Dps | IMEF | Encapsulin |
|---|---|---|---|---|---|
| [Bacillus] aminovorans DSM 1314 | no | no | OAH56278 | WP_063965031 | WP_063965030 |
| [Bacillus] aminovorans DSM 4337 | no | no | OAH62575 | OAH53583 | OAH53584 |
| Alteribacillus bidgolensis P4B | no | WP_091586138 | no | SDI49454 | SDI49426 |
| Alteribacillus iranensis DSM 23995 | no | WP_091663879 | WP_091663394 | WP_091658128 | WP_091658125 |
| Aneurinibacillus tyrosinisolvens | OIJ04211 | no | WP_071319460 | WP_047151354 | WP_047151353 |
| Aneurinibacillus migulanus DSM 2895 | WP_043066706 | WP_043065797 | WP_021619211 | WP_043068886 | WP_043068885 |
| Aneurinibacillus sp. XH2 | no | no | WP_057898153 | PGF_00166540 | WP_043068885 |
| Aneurinibacillus thermoaerophilus L 420-91 | no | no | WP_091260354 | SDH31038 | SDH31061 |
| Quasibacillus thermotolerans MTCC 8252 | no | no | WP_039234032 | WP_039238473 | WP_039238471 |
| Bacillus azotoformans LMG 9581 | no | no | WP_087946144 | EKN64196 | EKN64195 |
| Bacillus azotoformans MEV2011 | no | no | WP_035195831 | KEF38094 | KEF38093 |
| Bacillus methanolicus MGA3 | no | no | WP_003347246 | AIE58873 | AIE58874 |
| Bacillus methanolicus PB1 | no | no | WP_003351303 | WP_003351118 | WP_003351117 |
| Bacillus sp. 1NLA3E | no | no | WP_015595456 | WP_015593352 | WP_015593353 |
| Bacillus sp. FJAT-27238 | no | no | KMZ44901 | WP_057773495 | WP_057773497 |
| Bacillus sp. OK048 | no | no | WP_090761658 | WP_090761391 | WP_090761393 |
| Bacillus sp. OV166 | no | no | WP_088089380 | SMQ84107 | SMQ84105 |
| Bacillus sp. strain JF8 | no | no | AGT33197 | AGT31241 | AGT31240 |
| Bacillus thermotolerans SGZ-8 | no | no | WP_039234032 | QY97_0899 | WP_015593353 |
| Brevibacillus agri BAB-2500 | no | no | WP_081592058 | ELK43221 | ELK43220 |
| Brevibacillus borstelensis AK1 | no | no | WP_003388077 | WP_003389153 | WP_003389152 |
| Brevibacillus brevis NBRC 100599 | no | no | BAH42285 | WP_015892426 | WP_007721249 |
| Brevibacillus brevis ATCC 35690 | no | no | no | WP_016742089 | WP_007721249 |
| Brevibacillus brevis DZQ7 | no | no | WP_083261598 | WP_064202172 | WP_007721249 |
| Brevibacillus choshinensis DSM 8552 | no | no | WP_055747325 | WP_055744591 | WP_055744592 |
| Brevibacillus formosus DSM 9885 | no | no | WP_047070550 | WP_047071899 | WP_007721249 |
| Brevibacillus formosus NF2 | no | no | WP_088906278 | WP_047071899 | WP_007721249 |
| Brevibacillus laterosporus DSM 25 | no | no | WP_018671993 | WP_003334817 | WP_003334816 |

| | | | | | |
|---|---|---|---|---|---|
| Brevibacillus laterosporus GI-9 | no | no | WP_018671993 | WP_003334817 | WP_003334816 |
| Brevibacillus laterosporus LMG 15441 | no | no | WP_018671993 | WP_003334817 | WP_003334816 |
| Brevibacillus panacihumi W25 | no | no | WP_023556060 | WP_023557771 | WP_023557772 |
| Brevibacillus parabrevis CN1 | no | no | WP_083955677 | WP_122964006 | WP_063228109 |
| Brevibacillus reuszeri DSM 9887 | no | no | WP_103109718 | WP_049742502 | WP_049742503 |
| Brevibacillus sp. BC25 | no | no | WP_007726053 | WP_007721247 | WP_007721249 |
| Brevibacillus sp. CF112 | no | no | WP_007781621 | WP_007783174 | WP_007783173 |
| Brevibacillus sp. OK042 | no | no | WP_092266183 | WP_092268779 | WP_092268777 |
| Brevibacillus sp. SKDU10 | no | no | WP_082890860 | WP_064017301 | WP_003334816 |
| Brevibacillus sp. WF146 | no | no | WP_044898308 | WP_029098686 | WP_065067008 |
| Domibacillus antri | no | no | WP_075399286 | WP_075398991 | WP_075398990 |
| Domibacillus enclensis DSM 25145 | no | no | WP_045852489 | WP_045851598 | WP_045851597 |
| Domibacillus iocasae DSM 29979 | no | no | WP_069938694 | WP_069939583 | WP_069939582 |
| Geobacillus kaustophilus GBlys | no | no | WP_014196600 | WP_044731961 | WP_044731962 |
| Geobacillus kaustophilus Et2/3 | no | no | WP_044733039 | BAD75202 | BAD75201 |
| Geobacillus lituanicus N-3 | no | no | WP_033014567 | WP_047757677 | WP_100659874 |
| Geobacillus sp. 12AMOR1 | no | no | AKM20138 | AKM18226 | AKM18225 |
| Geobacillus sp. 15 | no | no | KZM53562 | KZM56073 | KZM56072 |
| Geobacillus sp. 46C-IIa | no | no | WP_081208050 | WP_081207017 | WP_081206634 |
| Geobacillus sp. A8 | no | no | WP_011232333 | WP_014195239 | WP_119877568 |
| Geobacillus sp. B4113_201601 | no | no | WP_033018292 | WP_033843367 | WP_119877568 |
| Geobacillus sp. CAMR5420 | no | no | KDE47147 | WP_033024921 | WP_033024920 |
| Geobacillus sp. LC300 | no | no | AKU26136 | AKU27458 | AKU27457 |
| Geobacillus sp. MAS1 | no | no | ESU73485 | ESU72279 | ESU72278 |
| Geobacillus sp. PA-3 | no | no | KQB92177 | KQB94129 | no |
| Geobacillus sp. Sah69 | no | no | KQC46690 | KQC48288 | KQC48287 |
| Geobacillus sp. Y412MC52 | no | no | ADU95335 | ADU93311 | ADU93310 |
| Geobacillus stearothermophilus 10 | no | no | ALA69799 | WP_013523228 | WP_013523227 |
| Geobacillus stearothermophilus strain 22 | no | no | OAO85981 | WP_049626374 | WP_049626373 |
| Geobacillus stearothermophilus strain 53 | no | no | WP_033014567 | WP_033024921 | WP_033024920 |
| Geobacillus stearothermophilus strain A1 | no | no | KMY59742 | KMY59381 | KMY59380 |
| Geobacillus stearothermophilus strain B4109 | no | no | WP_033014567 | WP_033024921 | WP_033024920 |
| Geobacillus stearothermophilus strain B4114 | no | no | WP_033014567 | WP_033024921 | WP_033024920 |
| Geobacillus subterraneus KCTC 3922 | no | no | WP_063164935 | AMX84296 | AMX84297 |
| Geobacillus thermocatenulatus BGSC 93A1 | no | no | WP_025950204 | WP_014195239 | WP_119877568 |
| Geobacillus thermodenitrificans T12 | no | no | WP_008880962 | EDY07466 | |
| Geobacillus thermoleovorans B23 | no | no | WP_011232333 | WP_014195239 | WP_011230417 |
| Geobacillus thermoleovorans CCB_US3_UF5 | no | no | AGE23447 | AEV18405 | AEV18404 |

| | | | | | |
|---|---|---|---|---|---|
| Geobacillus thermoleovorans strain FJAT-2391 | no | no | AKU26136 | AWO73557 | AWO73558 |
| Lihuaxuella thermophila strain DSM 46701 | no | WP_089964625 | no | WP_089972673 | WP_089972676 |
| Sporosarcina globispora DSM 4 | no | no | WP_053434019 | WP_053433170 | WP_053437549 |
| Thalassobacillus cyri CCM7597 | no | WP_093043674 | no | WP_093041210 | WP_093041212 |
| Thermoflavimicrobium dichotomicum DSM 44778 | no | no | no | WP_093230746 | WP_093230618 |
| **Absent in:** | **97%** | **93%** | **7%** | | |
| | **Ftn and Bfr missing: 92%** | | | | |

**Table S2. Cryo-EM data collection statistics for IMEF-loaded encapsulin.**

| **Cryo-EM data collection and processing** | **IMEF encapsulin** |
|---|---|
| Electron microscope | Tecnai F20 |
| Voltage (kV) | 200 |
| Electron dose ($e^-$/Å$^2$) | 44 |
| Physical pixel size (Å) | 1.28 |
| Number of collected movies | 601 |
| Defocus range (avg) (μm) | 1.0-3.0 (2.3) |
| Particle number for final map | 18,995 |
| Symmetry for final map | I (icosahedral) |
| Resolution (Å) | 3.85 |
| Map sharpening B-factor (Å$^2$) | -151 |
| **Atomic model refinement (ASU)** | |
| Number of chains in ASU | 7 (4 Enc, 3 TP) |
| Number of protein residues | 1130 |
| Number of atoms | 9064 |
| **Geometric parameters (r.m.s.d.)** | |
| Bond length (Å) | 0.007 |
| Bond angle (º) | 0.941 |
| **Ramachandran statistics** | |
| Residues favoured (%) | 87.4 |
| Residues allowed (%) | 12.6 |
| Residues disallowed (%) | 0.0 |
| Rotamer outliers (%) | 0.61 |
| Clashscore | 3.96 |

**Table S3. List of ferritin-like protein and IMEF protein IDs used to construct the phylogenetic tree shown in Figure 3A.** IMEF: Iron-mineralizing encapsulin-associated Firmicute cargo, EncFlp: ferritin-like proteins (Flps) found within encapsulin operons containing targeting peptides, noEncFlp: Flps found outside encapsulin operons not containing a targeting peptide, Bfr: bacterioferritin, Rr: rubrerythrin, Mam-Ftn: mammalian ferritin, Bac-Ftn: bacterial ferritin, Dps: DNA-binding proteins from starved cells.

| Flp-type | Protein ID |
|---|---|
| IMEF | WP_018672703 |
| IMEF | WP_005828111 |
| IMEF | WP_044897354 |
| IMEF | WP_003332271 |
| IMEF | WP_035178714 |
| IMEF | WP_028782218 |
| IMEF | WP_04180969 |
| IMEF | WP_031404939 |
| IMEF | WP_028401721 |
| IMEF | WP_003348284 |
| IMEF | WP_014195239 |
| IMEF | WP_045851598 |
| IMEF | WP_046179756 |
| IMEF | IMEF |
| IMEF | WP_045518068 |
| IMEF | WP_028401721 |
| IMEF | WP_003348284 |
| IMEF | WP_043068886 |
| IMEF | WP_047151354 |
| EncFlp | ACL70345 |
| EncFlp | CAE09437 |
| EncFlp | ABR50494 |
| EncFlp | TmFlp |
| EncFlp | CAN95442 |
| EncFlp | ACY16335 |
| EncFlp | CAJ63146 |
| EncFlp | RrFlp |
| noEncFlp | AJP49228 |
| noEncFlp | ACV25530 |
| noEncFlp | AHK79895 |
| noEncFlp | AEF99408 |
| noEncFlp | CCE22839 |
| noEncFlp | AFL74931 |
| noEncFlp | ACL71928 |
| noEncFlp | AAZ98418 |
| noEncFlp | AHF03132 |
| noEncFlp | AKH19188 |
| noEncFlp | AKH68756 |
| noEncFlp | AGS38469 |
| noEncFlp | AFZ34279 |
| noEncFlp | CAD84078 |
| noEncFlp | BAP56432 |
| noEncFlp | ACB49386 |
| EncFlp | ABF90650 |
| EncFlp | MxFlp |
| EncFlp | ADO71860 |
| EncFlp | ABF92698 |
| EncFlp | ABN52731 |
| EncFlp | ABS55612 |
| EncFlp | ABV34473 |
| EncFlp | AAL81316 |
| EncFlp | AFN03982 |

| | |
|---|---|
| Bfr | EAQ77221 |
| Bfr | BAF71400 |
| Bfr | CAJ12841 |
| Bfr | ABQ95053 |
| Bfr | ABR71746 |
| Bfr | ABC22995 |
| Bfr | BFR_ECOLI |
| Bfr | AKH68832 |
| Bfr | CEL31094 |
| Bfr | BFR_AZOVI |
| Bfr | KIL04563 |
| Bfr | EPP19171 |
| Bfr | KMN33871 |
| Bfr | AJQ92466 |
| Bfr | CUR48233 |
| Bfr | ABC24268 |
| Bfr | ABR86521 |
| Rr | AAB88944 |
| Rr | AAB85322 |
| Rr | ACM24105 |
| Rr | CEP79124 |
| Rr | AIG98643 |
| Rr | AEH24665 |
| Rr | Q9V0A0_PYRAB |
| Rr | KUH34502 |
| Rr | Q5JF11_PYRKO |
| Rr | 3MPS_A |
| Rr | AAB90407 |
| Rr | AAB90419 |
| Rr | RUBY_DESVH |
| Rr | EEG74317 |
| Rr | 1J30_A |
| Rr | 4DI0_A |
| Rr | EDL54893 |
| Rr | AAK86067 |
| Mam-Ftn | XP_018870100 |
| Mam-Ftn | FRIL_PONAB |
| Mam-Ftn | FRIL_HUMAN |
| Mam-Ftn | FRIL1_MOUSE |
| Mam-Ftn | NP_001266795 |
| Mam-Ftn | XP_005005114 |
| Mam-Ftn | BAG82928 |
| Mam-Ftn | ELR56618 |
| Mam-Ftn | AAH61303 |
| Mam-Ftn | FRI2_LITCT |
| Mam-Ftn | FRIM_SALSA |
| Mam-Ftn | FRIH_TRENE |
| Mam-Ftn | FTMT_HUMAN |
| Mam-Ftn | 2FHA |
| Mam-Ftn | FRIH_MOUSE |
| Bac-Ftn | 4ITW_A |
| Bac-Ftn | EAQ74833 |
| Bac-Ftn | BFRB_MYCTE |
| Bac-Ftn | ABQ03725 |
| Bac-Ftn | EDM65666 |
| Bac-Ftn | FTNA_ECOLI |
| Bac-Ftn | CAL34758 |
| Bac-Ftn | FTN_HELPJ |
| Bac-Ftn | ABN51258 |
| Bac-Ftn | 2JD7_A |
| Bac-Ftn | ABT93906 |
| Bac-Ftn | ADD56915 |

| | |
|---|---|
| Dps | DPS_CAMJE |
| Dps | ABC21138 |
| Dps | AEP02251 |
| Dps | KIO64845 |
| Dps | DPS_BREBE |
| Dps | KKX55325 |
| Dps | PCN43109 |
| Dps | KPC73845 |
| Dps | AHX18156 |
| Dps | KQL50044 |
| Dps | ACS25476 |
| Dps | OQP01613 |
| Dps | EQB94255 |
| Dps | WP_044733039 |
| Dps | PAY11715 |
| Dps | Q65FU7_BACLD |
| Dps | DP_STRSU |
| Dps | DPS_LISMO |
| Dps | EAQ79922 |
| Dps | DPS_MYCSM |
| Dps | DPS_AGRFC |
| Dps | DPS_ECOLI |
| Dps | ABC22300 |

**Table S4. X-ray structure determination and refinement statistics for the IMEF cargo protein.**

| | |
|---|---|
| PDB ID | 6N63 |
| Unit cell (Å)[a] | a = b = 81.4, c = 65.9 |
| Spacegroup | P $4_1$ $2_1$ 2 |
| Resolution range (Å)[a] | 58.0 – 1.72 (1.78 – 1.72) |
| Wavelength (Å) | 0.9762 |
| Observed reflections | 219,893 |
| Unique reflections | 24,131 |
| Completeness (%) | 99.7 (100.0) |
| Redundancy | 9.1 (8.1) |
| $R_{pim}$(%)[b] | 0.026 (0.504) |
| Overall $<I/\sigma(I)>$ | 29.0 (1.0) |
| $CC_{1/2}$ | 0.982 (0.681) |
| $R_{cryst}^{c}/R_{free}^{c}$ (%) | 19.4/22.0 |
| Ramachandran plot (%) | |
| Favored/allowed/outliers | 97.8 / 2.2 / 0.0 |
| Bond lengths[d] (Å) | 0.018 |
| Bond angles[d] (°) | 1.430 |
| Average B-factors (Å) | |
| Protein | 49.4 |
| Waters | 52.3 |
| Other | 40.5 |

[a]Values in parentheses are for the highest resolution shell.

$$^{b}R_{pim} = \sqrt{\frac{1}{n-1}}\frac{\sum|I-\langle I\rangle|}{\sum I}$$

Where $I$ is the observed integrated intensity, $<I>$ is the average integrated intensity obtained from multiple measurements, and the summation is over all observed reflections.

$$^{c}R_{cryst} = \frac{\sum\left||F_o|-k|F_c|\right|}{\sum F_o}$$

$F_o$ and $F_c$ are the observed and calculated structure factors, respectively, and $k$ is a scaling factor. The summation is over all measurements. [d]$R_{free}$ is calculated as $R_{cryst}$ using 5% of the reflections chosen randomly and omitted from the refinement calculations. Model stereochemistry was analyzed using MolProbity (*49*).

[e]Bond lengths and angles are root-mean-square deviations from ideal values.

**Table S5. EELS data of electron dense cores of purified IMEF encapsulins produced in *E. coli* under high iron conditions.** #Fe from EELS analysis values were extracted using Gatan software. Errors combine estimated statistical error in this measurement with known error for cross section. For density calculations, particles were approximate as spheres.

| Particle ID | Diameter (nm) | Diameter Standard deviation (nm) | # Fe atoms from EELS analysis | Error in # Fe atoms | Calculated density (# Fe atoms/nm$^3$) |
|---|---|---|---|---|---|
| 1 | 22.8 | 1.45 | 19070 | 3756.79 | 3.07 |
| 2 | 20.5 | 1.37 | 7539 | 1485.18 | 1.67 |
| 3 | 19.2 | 2.38 | 8286 | 1632.34 | 2.25 |
| 4 | 12 | 1.73 | 2512 | 494.86 | 2.78 |
| 5 | 16.9 | 2.57 | 2836 | 558.69 | 1.13 |
| 6 | 19.4 | 3.64 | 8171 | 1609.69 | 2.13 |
| 7 | 15.4 | 0.46 | 3137 | 617.99 | 1.64 |
| 8 | 20.8 | 0.64 | 7327 | 1443.42 | 1.55 |
| 9 | 20.7 | 0.78 | 6574 | 1295.08 | 1.41 |
| 10 | 16.5 | 1.95 | 3936 | 775.39 | 1.66 |
| 11 | 19.8 | 4.43 | 7250 | 1428.25 | 1.78 |
| 12 | 20.3 | 2.70 | 9526 | 1876.62 | 2.17 |
| 13 | 23.6 | 1.80 | 23293 | 4588.72 | 3.40 |
| 14 | 19.5 | 2.36 | 7785 | 1533.65 | 2.00 |
| 15 | 18.7 | 2.40 | 3425 | 674.725 | 1.01 |
| 16 | 21.1 | 1.29 | 6721 | 1324.04 | 1.38 |
| 17 | 20.5 | 1.98 | 10170 | 2003.49 | 2.24 |
| 18 | 21.4 | 3.68 | 9840 | 1938.48 | 1.91 |
| 19 | 24.7 | 2.74 | 10968 | 2160.70 | 1.39 |
| 20 | 25.1 | 2.35 | 14739 | 2903.58 | 1.77 |
| 21 | 28.6 | 2.19 | 13124 | 2585.43 | 1.07 |
| 22 | 22.23 | 1.37 | 6783 | 1336.25 | 1.17 |
| Average | | | | | 1.85 |
